## Supplementary Text and Figures for "Regional patterns of human cortex development correlate with underlying neurobiology"

Supplementary Text S1.1 to S1.5.

Supplementary Figures S1 to S28.

Supplementary References.

Separate files include:

Supplementary Data S1 to S5 are provided in a separate Excel file.

Supplementary Movies S1 and S2 are provided in separate GIF files.

**Abbreviations:**

| | |
|---|---|
| CT | cortical thickness |
| MRI | magnetic resonance imaging |
| mRNA | messenger ribonucleic acid |
| ABA | Allen brain atlas |
| MNI | Montreal Neurological Institute |
| FDR | false-discovery rate |
| ABCD | Adolescent Cognitive Brain and Development Study℠ (ABCD Study®) |
| ANCOVA | analysis of covariance |
| ni | nuclear imaging-derived brain atlas |
| ce | cell marker-derived brain atlas |
| mr | MRI-derived brain atlas |
| SV2A | synaptic vesicle glycoprotein 2A |
| M1 | muscarinic receptor 1 |
| mGluR5 | metabotropic glutamate receptor 5 |
| 5HT1a/1b/2a/4/6 | serotonin receptor 1a/2a/4/6 |
| CB | cannabinoid receptor 1 |
| GABAa | γ-aminobutyric acid receptor A |
| HDAC | histone deacetylase |
| 5HTT | serotonin transporter |
| FDOPA | fluorodopa |
| DAT | dopamine transporter |
| D1/2 | dopamine receptor 1/2 |
| NMDA | N-methyl-D-aspartate glutamate receptor |
| GI | glycolytic index |
| MU | mu opioid receptor |
| A4B2 | α4β2 nicotinic receptor |
| VAChT | vesicular acetylcholine transporter |
| NET | noradrenaline transporter |
| CBF | cerebral blood flow |
| CMRglu | cerebral metabolic rate of glucose |
| COX1 | cyclooxygenase 1 |
| H3 | histamine receptor 3 |
| TSPO | translocator protein |
| Ex | excitatory neurons |
| In | inhibitory neurons |
| Oligo | oligodendrocytes |
| Endo | endothelial cells |
| Micro | microglia |
| OPC | oligodendrocyte progenitor cells |
| Astro | astrocytes |

### 1. Supplementary Methods and Results

#### 1.1. Correlating human cortex development patterns with adult neurobiological markers

As laid out briefly in the main manuscript, our rationale is based on the following premises:

(i) **X** is a neurobiological entity (e.g., dendrites, specific neurons, or neurotransmitter receptors) that changes with human neurodevelopment and aging.

(ii) Changes in **X** might have downstream effects on CT as measured with MRI, either directly (e.g., more/less neurons or dendrites lead to higher/lower CT) or indirectly (e.g., higher density of a neurotransmitter receptor as a proxy of certain cellular structures developing/degrading; cortical myelination leading to changes in gray-white-matter contrast). Alternatively, there might be a common neurobiological process **Y**, which independently affects both **X** and CT, leading to a correlation between changes in **X** and changes in CT that does not require a direct causal relationship between **X** and CT but would still imply a common underlying mechanism shaping both.

(iii) **X** is in appearance and strength distributed non-uniformly across cortical regions, resulting in a (more or less) specific spatial distribution.

(iv) The cross-regional distribution of **X** is not necessarily stable across the human lifespan, but it might vary with neurodevelopment and aging. However, a major change of **X**'s distribution would require that multiple brain regions change their relative ranks of **X** density/expression/activity as compared to, e.g., a general decrease of **X** in all regions or a marginal increase of **X** in one region that does not lead to a relevant "region rank-increase". Such a major reorganization process is relatively more likely during neurodevelopment (i.e., pre-birth to maximally early adulthood) and aging (late adulthood) than during the relatively stable middle adulthood period.

(v) If changes in **X** or **Y** led to changes in CT, the spatial distribution of these CT changes would likely resemble the "steady state" adult spatial distribution of **X/Y**. This can be assumed as the stable adult distribution of a given neurobiological entity has to be the result of its neurodevelopment, and also the beginning-point of its aging processes. Therefore, changes in **X/Y** that lead to CT changes will likely be strongest in cortex regions with high density/expression/activity in **X/Y**'s "steady state" as this is either the state that they are approaching (neurodevelopmental CT changes) or coming from (aging CT changes).

From these assumptions follows that an observed spatial correlation between (i) the distribution of **X** as measured during the stable period and (ii) the distribution of CT changes during a given developmental period could have resulted from a developmental process that **X** is subject to. As noted in the main manuscript, this by no means is the only explanation for such CT changes, which is discussed in detail in the manuscript's discussion. Furthermore, although we consider them conceptually sound, it is currently not possible to prove all of our assumptions and specific research on the matter – especially on cross-regional spatial patterns of (developmental) neurobiological processes – is extremely scarce. Finally, the described framework assumes an idealized scenario in which neither measures of CT change nor of **X** are subject to noise and in which all "neurobiological markers" of **X** are measured during the stable mid-adulthood period.

#### 1.2. Molecular and cellular neurobiological markers

##### *1.2.1. Atlas sources*

As neurobiological markers of cell populations, processes, and systems potentially underlying CT changes, we collected 27 *in vivo* nuclear imaging atlases (20 neurotransmitter systems, cerebral glucose uptake, blood flow, aerobic glycolysis, synaptic density, transcriptomic activity, and two atlases capturing brain immune function), an MRI-derived atlas of cortical microstructure (T1w/T2w ratio), and 21 atlases of neuronal and glial cell types generated from Allen Brain Atlas mRNA expression data based on marker genes identified in adult human brain tissue (Supplementary Data S1; Fig. S1)[1–29].

##### *1.2.2. Processing of Allen Human Brain Atlas mRNA expression data*

Regional microarray expression data were obtained from 6 *postmortem* brains (1 female, age range 24.0–57.0 years, mean age 42.50 ± 13.38 years) provided by the Allen Human Brain Atlas (https://human.brain-map.org)[12]. Data were processed with the abagen toolbox (version 0.1.3; https://github.com/rmarkello/abagen)[30] using a 148-region surface-based atlas in fsaverage5 space[31].

First, microarray probes were reannotated using data provided by Arnatkevičiūtė et al.[32]; probes not matched to a valid Entrez ID were discarded. Next, probes were filtered based on their expression intensity relative to background noise[33], such that probes with intensity less than the background in ≥ 50% of samples across donors were discarded, yielding 31,569 probes. When multiple probes indexed the expression of the same gene, we selected and used the probe with the most consistent pattern of regional variation across donors (i.e., differential stability[34]), calculated with:

$$\Delta_S(p) = \frac{1}{\binom{N}{2}} \sum_{i=1}^{N-1} \sum_{j=i+1}^{N} \rho\left[B_i(p), B_j(p)\right]$$

where $p$ is Spearman's rank correlation of the expression of a single probe, p, across regions in two donors $B_i$ and $B_j$, and N is the total number of donors. Here, regions correspond to the structural designations provided in the ontology from the Allen Human Brain Atlas. The MNI coordinates of tissue samples were updated to those generated via non-linear registration using the Advanced Normalization Tools (ANTs; https://github.com/chrisfilo/alleninf). To increase spatial coverage, tissue samples were mirrored bilaterally across the left and right hemispheres. Samples were assigned to brain regions by minimizing the Euclidean distance between the MNI coordinates of each sample and the nearest surface vertex. Samples where the Euclidean distance to the nearest vertex was more than 2 standard deviations above the mean distance for all samples belonging to that donor were excluded. To reduce the potential for misassignment, sample-to-region matching was constrained by hemisphere and gross structural divisions (i.e., cortex, subcortex/brainstem, and cerebellum, such that e.g., a sample in the left cortex could only be assigned to an atlas parcel in the left cortex[32]). All tissue samples not assigned to a brain region in the provided atlas were discarded. Inter-subject variation was addressed by normalizing tissue sample expression values across genes using a robust sigmoid function[35]:

$$x_{\text{norm}} = \frac{1}{1 + \exp - \left(\frac{(x - \langle x \rangle)}{\text{IQR}_x}\right)}$$

where $\langle x \rangle$ is the median and $IQR_x$ is the normalized interquartile range of the expression of a single tissue sample across genes. Normalized expression values were then rescaled to the unit interval:

$$x_{\text{scaled}} = \frac{x_{\text{norm}} - \min(x_{\text{norm}})}{\max(x_{\text{norm}}) - \min(x_{\text{norm}})}$$

Gene expression values were then normalized across tissue samples using an identical procedure. Samples assigned to the same brain region were averaged separately for each donor and then across donors, yielding a regional expression matrix with 148 rows, corresponding to brain regions, and 15,633 columns, corresponding to the retained genes.

##### *1.2.3. Temporal stability of original brain atlases*

Our analyses make use of the spatial pattern present in every given brain map to draw relationships between brain maps from different sources and levels of biological organization. This pattern is encoded in the relative ranks of the metric in question across the cortex regions. To retain good interpretability of our findings, the multimodal atlases should, in the best case, show a high stability of these ranks over the adult lifespan. We tested for this by collecting the available PET maps derived from the same tracer in healthy adult samples which's mean age different from the atlases used in our main analyses[11,16,17,36–38].

Spearman correlations across cortex regions indicated a high stability ($r \geq .77$) which, based on the limited data available, did not depend on sample age (Fig. S2).

##### *1.2.4. Factor analyses validation against null data*

In sensitivity analyses, we tested if the factor solutions estimated on the original brain atlases explained significantly more variance in the original data than factor solutions estimated on permuted brain atlases. Using the same autocorrelation-preserving method as in our remaining analyses[39], we generated $n = 10{,}000$ permuted brain maps. Each of these null datasets was z-standardized and used to fit a factor analysis (same settings as used in the main analyses) with $n = 10$ factors, separately for nuclear imaging and cell types. Explained variance in the observed data was calculated as the fraction of (i) the variance in the observed dataset and (ii) the variance in each null dataset. Based on the resulting null distributions of explained variance scores, two empirical $p$ values were calculated for the nuclear imaging and the cell type datasets.

At an alpha level of $p > .05$, the "observed" factor analyses explained more variance in the observed data as compared to factor analyses estimated on null data (nuclear imaging: $p = .0317$; cell types: $p = .0495$; Fig. S3).

#### 1.3. Discovery analyses based on the Braincharts model

##### *1.3.1. Modeled CT data*

Modeled CT values across the lifespan for 148 cortex regions were extracted from the *Braincharts* normative model published by Rutherford et al.[40] (5–90 years with 0.5-year steps; separate female and male data from approximately 58,000 subjects; $1^{st}$, $5^{th}$, $25^{th}$, $50^{th}$, $75^{th}$, $95^{th}$, and $99^{th}$ model percentile, age distribution: Fig. S5). As expected, lifespan cortical thickness development showed a general trend towards cortical thinning, with different trajectories across brain regions. While absolute values differed slightly by sex, the general trajectories and relative development were highly similar, so that we decided to average the data across sexes for all main analyses (Fig. S6A, Movie S1).

Cross-sectional colocalization analyses were based on the modeled CT data at each extracted timepoint. For analyses aiming to explain modeled longitudinal CT change patterns, we calculated timepoint-to-timepoint relative CT change of $50^{th}$ percentile data within a sliding window approach (5–90-year range, 5-year length, and 1-year steps). During childhood, adolescence, and up to young adulthood, precentral and temporal gyri showed the strongest relative increase of modeled CT while the remaining cortex showed thinning patterns. Generally, the strongest modeled CT changes across the lifespan (mostly thinning) occurred in the first and last third of life (Fig. S6B–C).

##### *1.3.2. Replication of explained CT change patterns using the Desikan-Killiany parcellation*

All analyses in this work are based on the 148-parcel (74 per hemisphere) Destrieux parcellation[31], subdividing the cortex into gyri and sulci. However, the 68-parcel (34 per hemisphere) Desikan-Killiany parcellation[41] (DK), which subdivides the cortex into gyri only, is more prevalent in the literature and is set as standard in FreeSurfer. We clearly argue that our approach requires a finer parcellation and therefore decided for the 148-parcel version. Furthermore, the Braincharts model is available for the Destrieux parcellation only. To nevertheless provide preliminary results on the basis of the DK parcellation, we transformed all surface data, i.e., both modeled CT change maps and the factor-level neurobiological markers from Destrieux to DK by (i) projecting the Destrieux data to the fsaverage5 surface and (ii) "re-parcellating" it using the DK parcels. While this is clearly not the most accurate approach, we argue that it is sufficient to approximate whether our results replicate with the DK parcellation, especially considering that many DK gyri will mostly be a combination of Destrieux gyri and sulci. Using this transformed data (Fig. S11A and B), we recalculated the first step of our regression analysis workflow (c.f., Figs. 3 and 4).

Using the DK parcellation, the overall explained modeled CT change pattern remained consistent with the Destrieux-based results (Fig. S11C and D). Most notably, $R^2$ values were overall considerably higher in the DK analyses regarding both univariate regression results (ranging up to $R^2$ = 70% for individual markers, e.g., *ni9-D2*) and multivariate regression results (approaching $R^2$ = 90%). As also the null values (grey shades in Fig. S11C and D) were distributed around a much higher range (median at about $R^2$ = 30% for univariate analyses), it appeared that this increase in explained variance was largely due to overparameterization or low degrees of freedom, respectively, in the regression models. These results speak to our focus on the more fine-grained parcellation.

##### *1.3.3. Potential confounding effects of Braincharts and neurobiological marker cohort ages*

We performed exploratory correlation analyses to exclude potential confounds brought about by the age distributions of the independent cohorts in regard to the explained modeled CT change patterns. First, to exclude potential biases due to model precision and dominance of single original cohorts, we tested if the explained modeled CT change across time windows was correlated to the number of subjects that went into model estimation per time window. A concerning case would have been negative correlations, pointing to high explained modeled CT change in time windows in which only few subjects went into model estimation (i.e., childhood and very late adulthood, see Fig. S5). Second, to exclude potential bias introduced by the average age of each neurobiological marker's source cohort, we correlated explained modeled CT change with the distance in years to the approximate average cohort age for each neurobiological marker (Data S1). Here, positive correlation would have been most concerning, possibly indicating cases in which, e.g., high explained modeled CT change in adolescence and aging was a result of using marker templates from the intermediate early and mid-adulthood age period. We performed these analyses with the univariate marker-wise explained modeled CT change (21 markers) as well as the multivariate explained CT change based on (i) only neuroimaging markers, (ii) only cell markers, and (iii) all combined markers. For the second analysis, we calculated the distance between each time window and each source age associated with the neurobiological markers by taking the mean of each time window [e.g., 7.5 for $\Delta(5,10)$ years] for all time windows, and calculating the absolute distance between each time window and the marker's age for a given marker (e.g., if marker age is 40 years: $|7.5 - 40| = 32.5$ years). As we worked with the factor level markers, we approximated factor-level marker ages by scaling the factor loadings for each marker to (0,1) and calculating, for each factor-level marker, the weighted mean of all original markers. Explained modeled CT change and age-related vectors were correlated using Spearman correlations and exact $p$ values were calculated based the explained modeled CT change null distributions already calculated for the main analyses (n = 10,000).

We did not find evidence for the outlined confounds of Braincharts and neurobiological marker source cohorts ages on explained modeled CT change. For the Braincharts age analysis, we observed only one significant negative correlation (uncorrected) for a neurobiological marker (ce6) that did not turn out relevant for our following analyses (Fig. S12). Regarding the neurobiological marker age analysis, we only found significant negative correlations for the combined molecular markers, *ni1,* and *ni6* (uncorrected) but no significant positive correlations (see above) and no systematic association pattern (Fig. S13).

##### *1.3.4. Brain-regional contributions to CT association patterns*

We evaluated brain-regional contributions to the overall explained modeled CT change by calculating the residual difference for each brain atlas as the difference in prediction errors resulting from a multivariate regression with and without the brain atlas included as predictor.

Generally, medial occipital, medial temporal, sensorimotor, and cingulate cortices influenced the explained modeled CT change patterns strongly. For *ce9-In8*, the premotor cortex, cuneus, and multiple frontopolar sulci were the most influential regions. *ce3-Micro-OPC* showed a similar pattern, while *ni9-D2* showed the strongest residual differences in the middle cingulate cortex, precuneus, insula, and temporal pole. The two metabolism factors (*ni4* and *ni6*) showed less pronounced patterns with generally occipitotemporal regions having the strongest influences. The two neurobiological markers relevant for midlife modeled CT change, *ni3-FDOPA-DAT-D1-NMDA and ni5-VAChT-NET*, again displayed a high relevance of lateral and medial somatosensory cortices as well as the precuneus, middle to anterior cingulate, and medial temporal regions. One marker associated to late adulthood modeled CT aging patterns, *ce4-In3-In2-Astro*, displayed the strongest influences by middle and anterior cingulate cortices and medial motor areas. Fig. 6 shows the respective patterns at each marker's maximum explained modeled CT change, Movie S2 illustrates how the observed patterns develop longitudinally, and Fig. S11 provides an overview for the complete lifespan.

##### *1.3.5. Evaluation of original in comparison to dimensionality-reduced neurobiological markers*

To demonstrate that the factor-level atlases were appropriately representing the original neurobiological brain atlases, we performed additional sets of (i) dominance analyses and (ii) univariate linear regressions for each factor-level atlas, using as predictors the 5 original atlases with the highest factor loadings if the absolute loading exceeded 0.3. FDR correction was performed across (i) all dominance analyses and (ii) all individual univariate regression separately.

All factors explained modeled CT change significantly (nominal $p < 0.05$). In all cases, the total explained modeled CT change $R^2$ peaks arising from each original atlas set occurred at the same time in life as observed for the factor-level atlases. For some factor-level atlases, we discovered that their peak contribution to explained modeled CT change was driven by a certain original atlas, while other original atlases were of lesser relevance. For *ni3-FDOPA-DAT-D1-NMDA*, we showed that the midlife peak was mostly driven by NMDA and DAT. Other strongly loading atlases – DOPA, D1, and NET – explained more modeled CT change before 25 and after 50 years. The contribution to early explained modeled CT change of *ni4-GI-5HT1b-MU-A4B2* was driven by GI, capturing aerobic glycolysis. For *ni5-VAChT-NET*, VAChT indeed accounted for most of the modeled CT change explained by the factor-level atlas during midlife. Furthermore, the A4B2 nicotinic receptor contributed to this factor. The relevance of *ni9-D2* for early explained modeled CT change was indeed driven by the D2 receptor, however the D1 receptor additionally loaded on the factor and accounted for around 15% of explained early modeled CT change. For *ce3-Micro-OPC*, the microglia distribution contributed more strongly, although explaining less modeled CT change in comparison to other predictors. Finally, *ce5-In6-Ex2* was dominated by Ex2 (layer 3/4 granule neurons), and we observed an additional contribution of Ex3 (layer 4 granule neurons) to *ce9-In8*. Fig. S14 illustrates detailed results, Fig. S15 shows the peak region-wise residual differences for each original atlas.

#### 1.4. Validation analyses based on ABCD and IMAGEN single-subject data

##### *1.4.1. ABCD and IMAGEN cohort demographics and quality control*

We obtained single-subject CT data from the ABCD Study®[42] and the IMAGEN[43] cohort studies to validate our findings. Data quality was ensured based on the manual ratings included in the ABCD dataset and on FreeSurfer's "number of surface defects" metric. Data S1 lists age and sex distributions (self-reported sex) for both cohorts and each timepoint. Fig. S18 shows the quality control metric distributions.

Regarding the ABCD dataset, initially, baseline data for 11,760 subjects and 2-year follow-up data for 7,829 subjects was available. After dropping one study site without longitudinal data and subjects with missing CT data, 11,716 and 7,818 datasets were retained. Subjects with low data quality were excluded, leading to 10,697 and 6,789 subjects. Concerning the IMAGEN dataset, 4,990 observations from 2,158 subjects were initially available. For 3,975 observations from 1,528 subjects, structural MRI data was available and successfully FreeSurfer-processed. After exclusion of subjects because of low data quality, 3,732 observations from 1,522 subjects were retained. From these 1,522 subjects, 1,412 had longitudinal data available.

Main analyses on these observed ABCD (T0–T2: n = 6,789) and IMAGEN (overall n = 1,412; T0–T8: n = 951; T0–T5: n = 1,142; T5–T8: n = 915) longitudinal datasets were performed independently from the Braincharts CT model after removing site-effects using ComBat-GAM[44]. For all analyses including subject-level predictions by the Braincharts model, the data was projected into the Braincharts model using healthy subsamples of both dataset's baseline data (n = 20 per site, 50% female, distributed across baseline age ranges) as "adaptation" cohorts according to Rutherford et al.[40]. For this purpose, "healthy" was defined as: no psychiatric diagnoses according to KSADS-parent (ABCD) or DAWBA (IMAGEN); no medical diagnoses according to "abcd_mx01.txt" (ABCD-only), and no history of traumatic brain injury (ABCD-only). As this adaptation cohort was then dropped from further analyses, sample sizes reduced for the IMAGEN cohort (overall n = 1,252; T0–T8: n = 842; T0–T5: n = 1,007; T5–T8: n = 826). This was not the case for the ABCD cohort as, here, baseline-only subjects were used for adaptation.

##### *1.4.2. Generalizability of individual CT prediction models*

We asked if the prediction of individual CT change patterns was generalizable from the median predictions of the normative model to the observed single subject data, i.e., we asked if a "one-size-fits-all" approach would have performed equally well. This was done by applying the parameters of the regression models estimated to predict each subject's *normative* CT change patterns to the same subject's *observed* change patterns, calculating the Pearson correlation between observed and predicted CT change patterns, and comparing these model fit metrics between the different models. To estimate the effect size, results were contrasted to null analyses in which each regression model was estimated using 1,000 permuted neurobiological marker maps.

While these one-size-fits-all models generally exceeded the predictive performance of permuted null models, they did not provide good fit for many individuals, thus highlighting the value of our individual differences-focused approach (Fig. S24).

##### *1.4.3. Effects of sex and site on explained CT change*

After having assessed how CT development on the single-subject level was explained from neurobiological markers, we evaluated whether the amount to which it was explained, varied with sex and study site.

ANCOVA models corrected for follow-up duration and site showed significantly more explained CT change in males as compared to females only in the IMAGEN dataset (timespans T0–T5 and T0–T8), but not in the ABCD data. Furthermore, we observed effects of site on explained CT change in ANCOVA models corrected for sex and follow-up duration in both datasets (all timespans). Note that this was done albite successful site-harmonization of the cross-sectional CT data (Data S3): For ABCD, ComBat-GAM harmonization removed site effects for all Destrieux parcels. For IMAGEN, harmonization reduced differences between sites but ANCOVAs controlled for sex, intracranial volume, and age still revealed significant differences for many parcels. For CT change, site effects were only slightly reduced due to harmonization of the underlying cross-sectional CT data and could be observed for most parcels (Data S4). Given that the main goal of the current analysis was to establish the feasibility of capturing associations between CT development and neurobiological markers on the individual level, clarification of the sources of sex- and site-effects will be a task for future investigations. Given the pattern of results, we assume that the observed differences by site, which notably grow stronger when looking at CT *change*, might not only be due to scanner variance, but possibly also biological, social, or behavioral differences, leading to diverging individual cortex development. Fig. S26 visualizes the observed group differences, Tabs. S3 and S4 show ANCOVA results.

##### *1.4.4. Effects of reference model predictive performance, subject-level deviations, and data quality on explained CT change*

We conducted correlational analyses to provide general indications of which factors influenced the extents to which CT change was explained in single-subject data.

First, CT change patterns of subjects who had more "normative" CT patterns at *baseline* (i.e., stronger correlation between observed and Braincharts-predicted baseline CT) were not consistently better explained. In contrast, subjects whose CT change patterns were more in line with the *change* patterns predicted by the Braincharts model showed higher explained CT change. Similarly, while we did not observe consistent relationships between metrics capturing how a subject deviated from the model-predictions (count of deviation Z scores > 2 and average absolute Z scores) and explained CT change, the longitudinal change in deviation metrics showed an association in all IMAGEN timespans: As expected, most subjects did not show strong changes in their deviations between timepoints, but subjects who showed *more or stronger deviations* at follow-up as compared to baseline tended towards *more explained CT change*. Whether such patterns could represent a potentially pathophysiological involvement of a certain neurobiological marker in neurodevelopment remains to be investigated. Finally, we observed that less CT change was explained in subjects with more surface defects (i.e., worse reconstruction quality) at follow-up (ABCD and IMAGEN) or at baseline (ABCD only). Fig. S27 shows the reported association patterns and provides Spearman correlation statistics.

#### 1.5. Software

Multimodal brain atlases were retrieved and processed from/with *neuromaps (0.0.2)*[21], *abagen (0.1.3)*[30], *JuSpace (1.3)*[45], or author sources. Analyses of associations between CT and cortical atlases were conducted using *JuSpyce 0.0.2*[46] in a *Python 3.9.11* environment[46]. *JuSpyce* (https://github.com/LeonDLotter/JuSpyce) is a toolbox allowing for flexible assessment and significance testing of associations between multimodal neuroimaging data, relying on imaging space transformations from *neuromaps*[21], brain surrogate map generation from *BrainSMASH (0.11.0)*[39], and several routines from *Nilearn (0.10.2)*[47], *scipy (1.12.0)*[48], *NiMARE (0.0.11)*[49], *statsmodels (0.14.1)*, *pingouin (0.5.4)*, *numpy (1.22.4)*, and *pandas (1.5.3)*. Visualizations were created using *matplotlib (3.8.3)*[50], *seaborn (0.11.0)*[51], and *surfplot (0.1.0)*[52]. The *PCNtoolkit (0.29.post1)*[53,54] was used to generate modeled CT data, as well as predicted CT data and deviation scores for ABCD and IMAGEN subjects; *neuroHarmonize (2.3.1)*[44] was used to independently harmonize the CT data across sites.

#### 2. Supplementary Figures

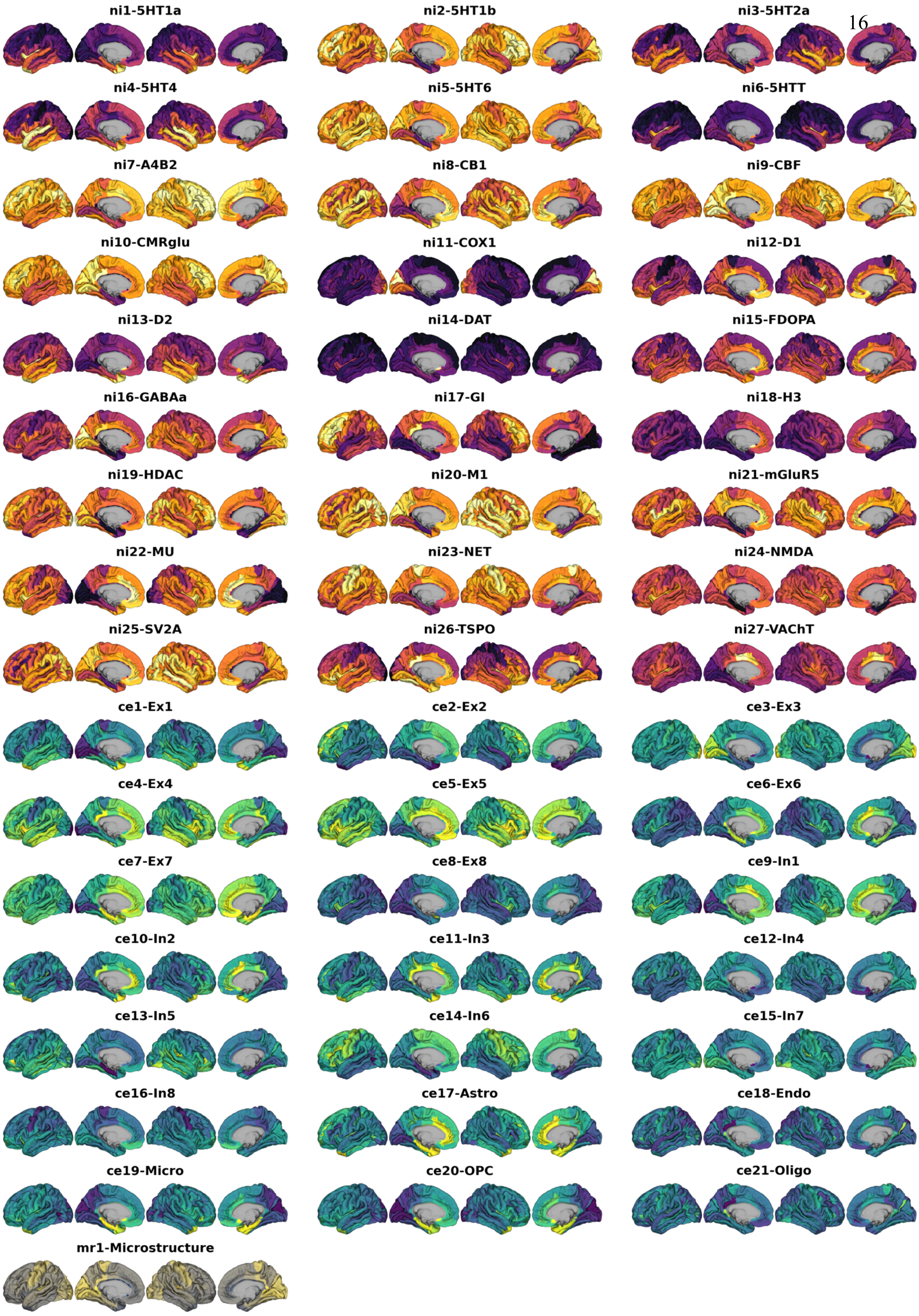
ni1-5HT1a
ni2-5HT1b
ni3-5HT2a
ni4-5HT4
ni5-5HT6
ni6-5HTT
ni7-A4B2
ni8-CB1
ni9-CBF
ni10-CMRglu
ni11-COX1
ni12-D1
ni13-D2
ni14-DAT
ni15-FDOPA
ni16-GABAa
ni17-GI
ni18-H3
ni19-HDAC
ni20-M1
ni21-mGluR5
ni22-MU
ni23-NET
ni24-NMDA
ni25-SV2A
ni26-TSPO
ni27-VAChT
ce1-Ex1
ce2-Ex2
ce3-Ex3
ce4-Ex4
ce5-Ex5
ce6-Ex6
ce7-Ex7
ce8-Ex8
ce9-In1
ce10-In2
ce11-In3
ce12-In4
ce13-In5
ce14-In6
ce15-In7
ce16-In8
ce17-Astro
ce18-Endo
ce19-Micro
ce20-OPC
ce21-Oligo
mr1-Microstructure

**Fig. S1: Multimodal nuclear imaging and neural cell type atlases**

Multimodal atlases after transformation to FreeSurfer space and parcellation into 148 cortical parcels (Destrieux parcellation[31]); **yellow-violet**: nuclear imaging markers, **yellow-green**: gene-expression, **yellow-gray**: microstructural; **yellow = higher density**. See Data S1 for individual descriptions and sources. Source data are provided as a Source Data file.

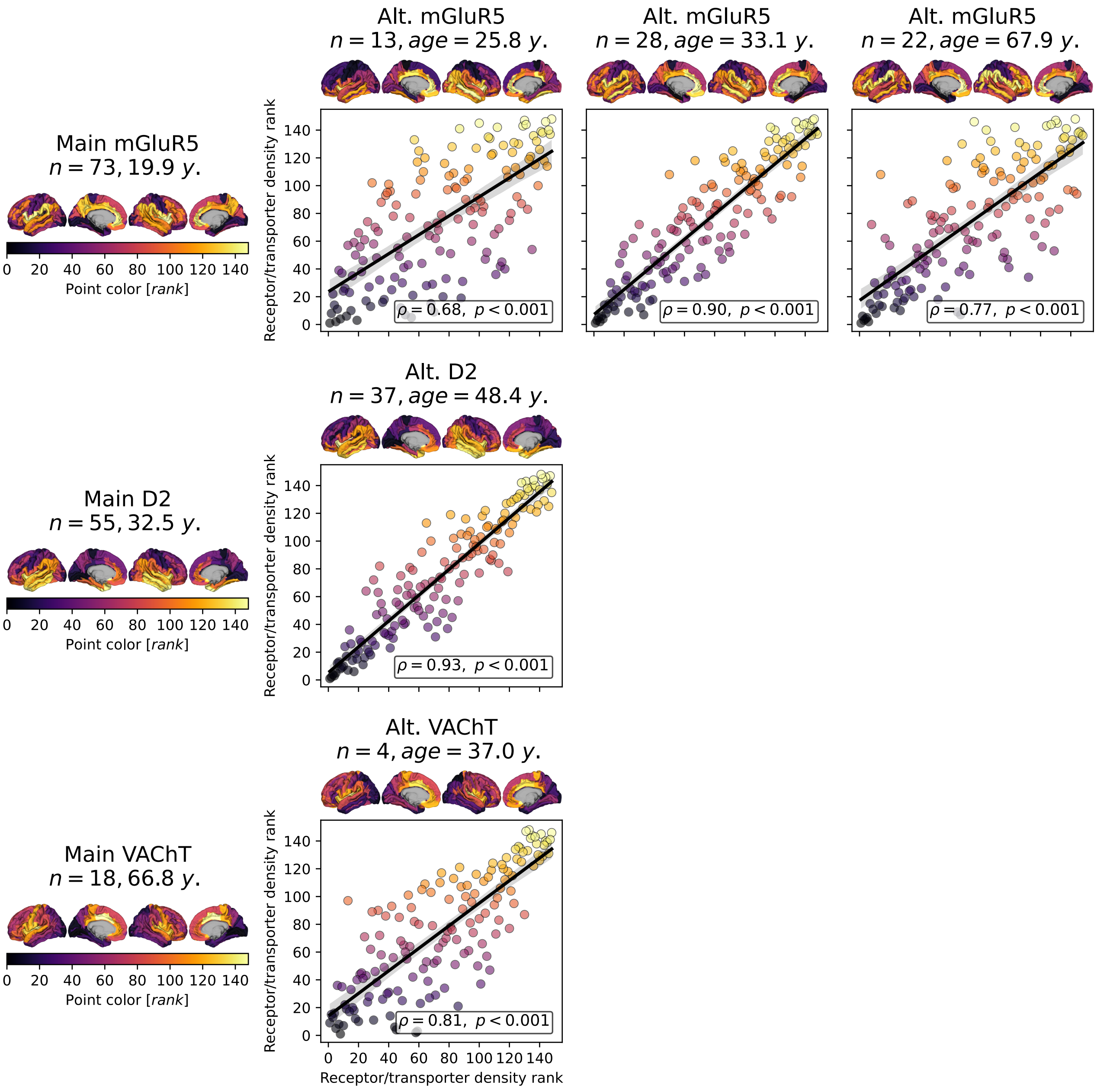

#### Fig. S2: Stability of regional PET atlas ranks throughout the adult lifespan

Comparison of PET atlases derived from independent healthy adult samples. We included all available PET atlases that used the same tracer but differed in regards to mean sample age, which resulted in three tracers for the glutamatergic, dopaminergic, and cholinergic systems. This analysis aimed to demonstrate, using the limited data available, that the *ranks* of regional tracer density across the cortex are stable during the adult lifespan. **Left side and y-axes**: Atlases used in the main analyses of this study[1,25,26]. **Scatter points** are colored according to the regional values. **Surface plots on top of each panel and x-axes**: Alternative atlases[11,16,17,36–38]. **Statistics**: Spearman correlations and parametric $p$ values. **Abbreviations**: Alt. = alternative, y. = years, mGluR5 = metabotropic glutamate receptor 5, D2 = dopamine receptor 2, VAChT = vesicular acetylcholine receptor. Source data are provided as a Source Data file.

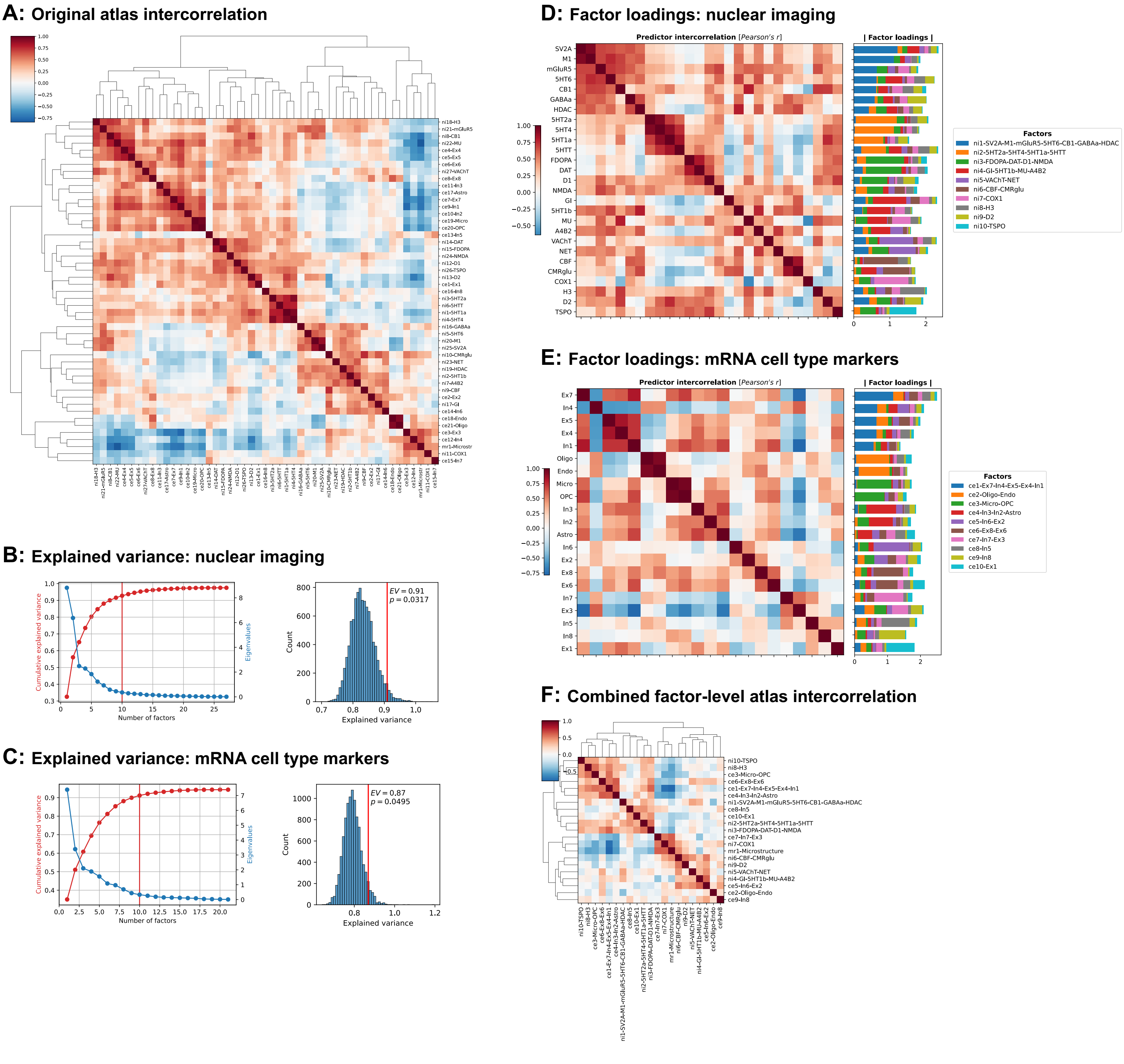

#### Fig. S3: Dimensionality reduction of multimodal atlases

**(A)** Spearman correlation matrix of original multimodal atlases. **(B) & (C)**: **Left**: Cumulative explained variance (red) and eigenvalues (blue) of unrotated factors in a minimum residual factor analysis on the nuclear imaging atlases **(B)** and mRNA expression atlases **(C)**. The red vertical line marks the threshold of factors explaining at least 1% of variance. **Right**: Null distribution of total variance explained scores, if factor analyses estimated on permuted brain maps ($n$ = 10,000) are used to explain variance in the original data. Red lines indicate the observed explained variance. **(D) & (E)**: Factors extracted from the nuclear imaging **(D)** and mRNA expression **(E)** datasets after promax rotation. The **heatmaps** show Pearson correlations between the original atlases before factor analysis. **Stacked bar plots** show factor loadings for each original atlas on each factor. Factor names were derived from assigning each original atlas to the factor it loaded on most so that each original atlas appears exactly once in the overall factor names. **(F)** Spearman correlation matrix of the 20 derived factors and the microstructural atlas. Source data are provided as a Source Data file.

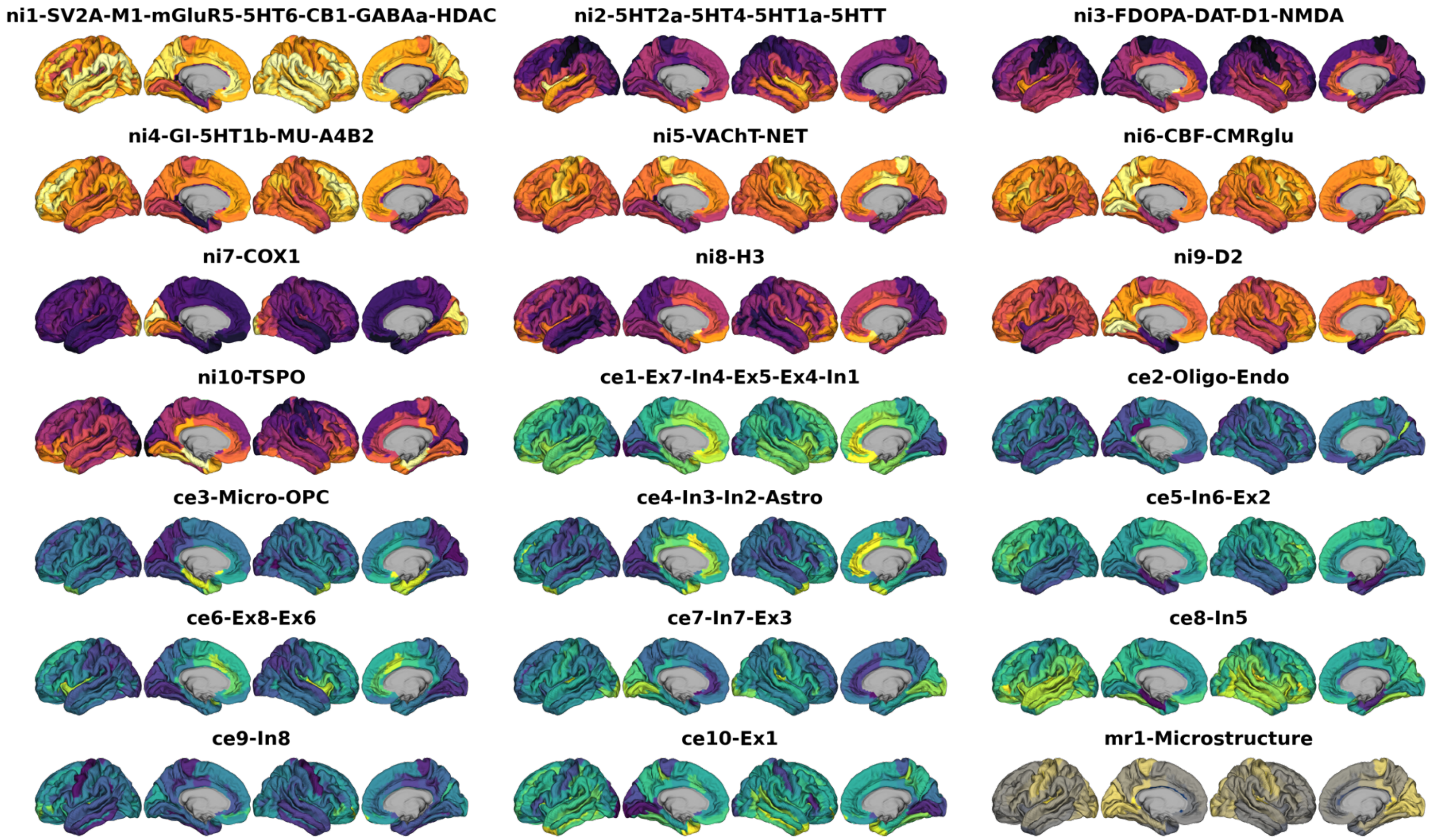

**Fig. S4: Neurobiological markers after factor analysis**

Parcellated brain atlases after dimensionality reduction, mapped to the cortex; **yellow-violet**: nuclear imaging markers, **yellow-green**: gene-expression, **yellow-gray**: microstructural; **yellow = higher density**. Source data are provided as a Source Data file.

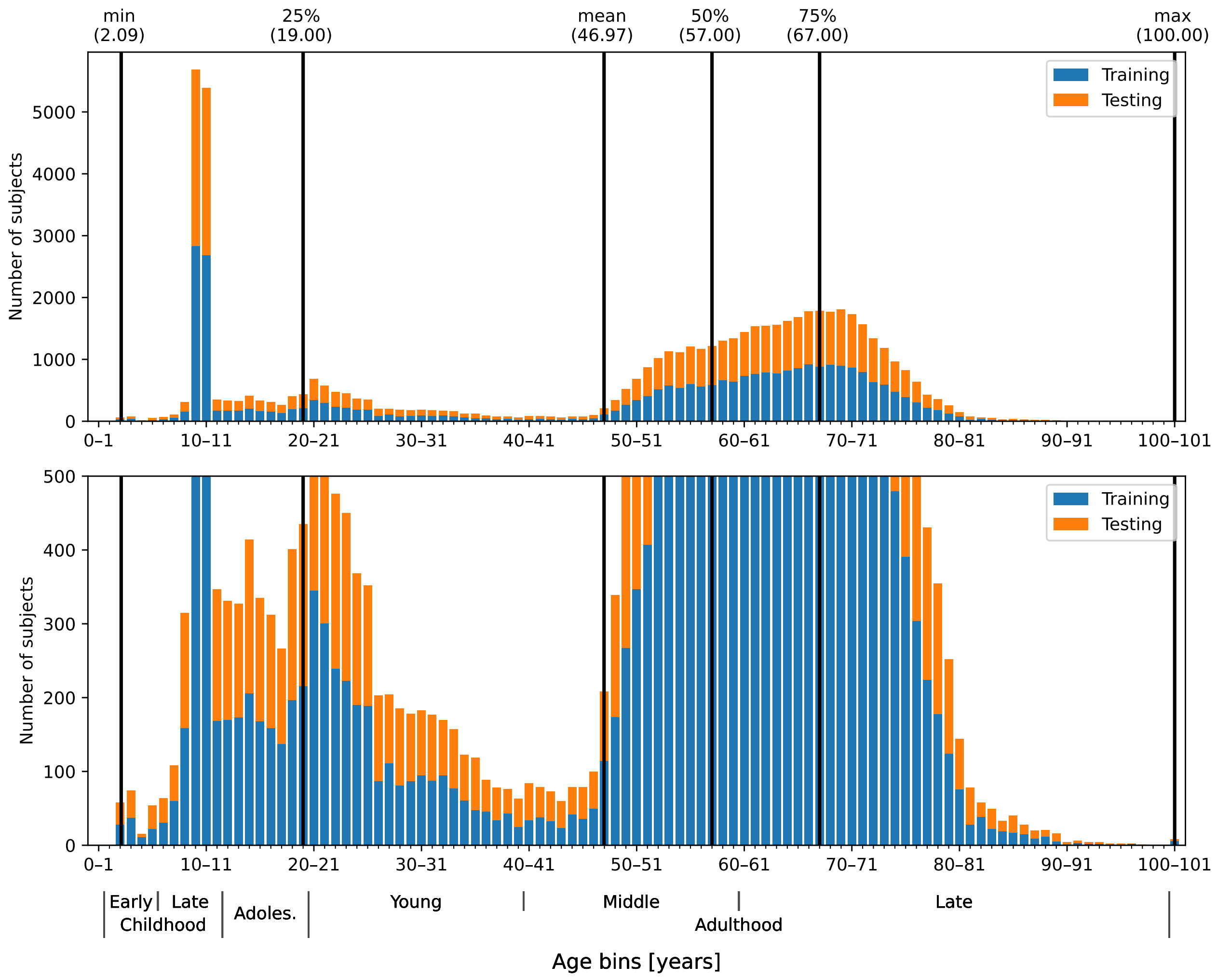

**Fig. S5: Age distributions of the Braincharts cohort**

Age distribution in Braincharts model, displayed as **stacked histograms** (**blue**: training data, **orange**: testing data) with 1-year bins (bin “0–1”: ≥ 0 to < 1 years, bin “1–2”: ≥ 1 to < 2 years, …). The lower panel is a copy of the upper panel with scaled y axis to visualize smaller bins. **Black vertical lines** show descriptive statistics for the whole dataset. The sharp peak at 9 to 11 years is caused by the ABCD study dataset, the smoother peak at 50 to 75 years is related to the UK Biobank study. Source data are provided as a Source Data file.

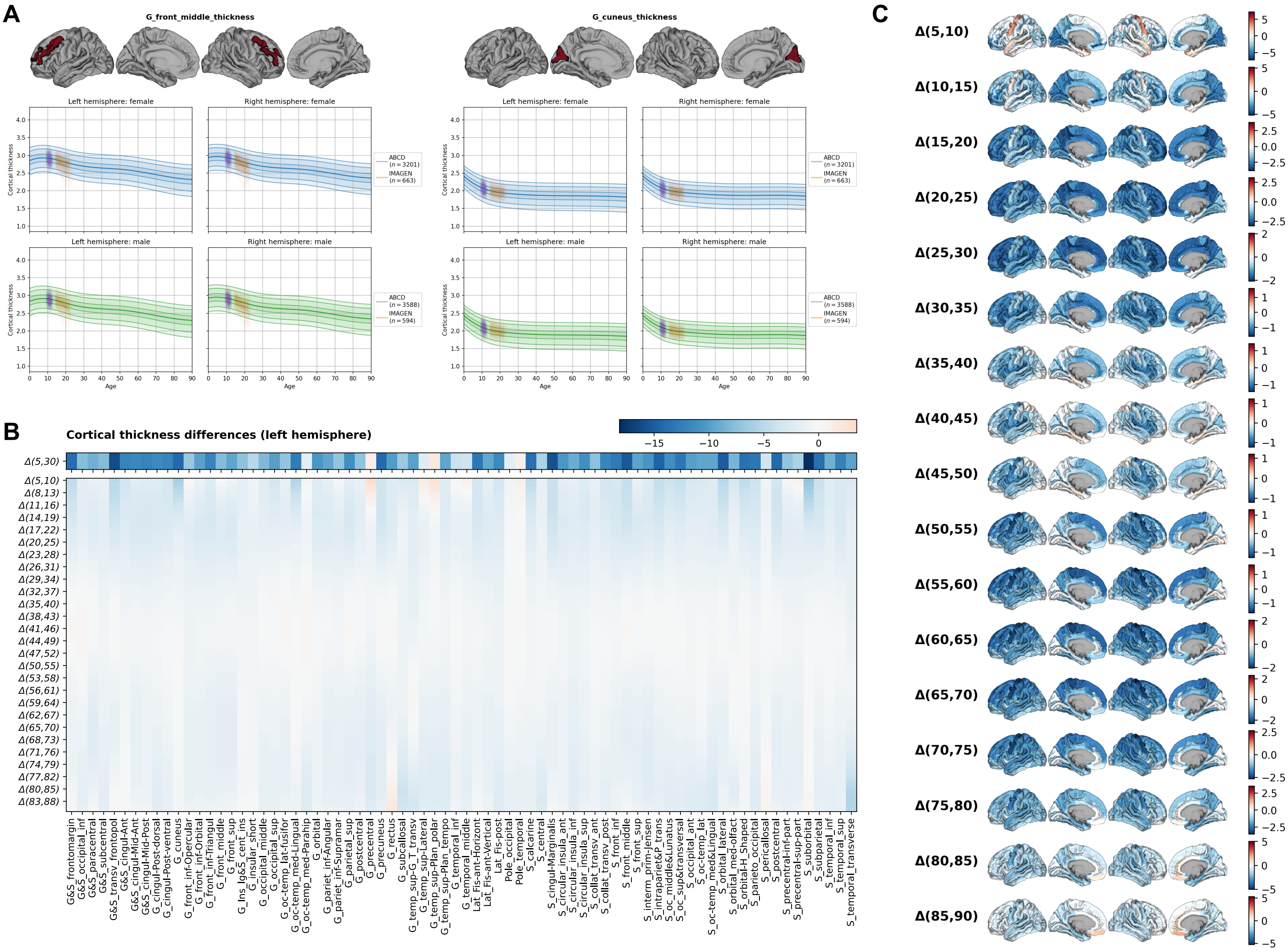

#### Fig. S6: Modeled lifespan cortical thickness development

Modeled cortical thickness data extracted from the reference model by Rutherford et al.[40], as well as ABCD[42] and IMAGEN cohorts[43]. **(A)** Exemplary modeled developmental trajectories of cortical thickness of two single bilateral brain regions for females (blue) and males (green). The **thickest line** represents the median, the **lighter lines** show the .01$^{th}$, .05$^{th}$, .25$^{th}$ as well as the .75$^{th}$, .95$^{th}$, and .99$^{th}$ centiles. The **clouds of lines** show ABCD and IMAGEN subjects after adaptation to the Braincharts model at each study time point. See Movie S1 for all brain regions. **(B)** Left-hemispheric region-wise relative cortical thickness differences in percent-change from 5 to 30 years (upper row) and from 5 to 90 years, estimated using a sliding window with 1-year steps and 5-year window length. **(C)** Modeled cortical thickness development in 5-year steps, plotted are region-wise percent-change values. Source data are provided as a Source Data file.

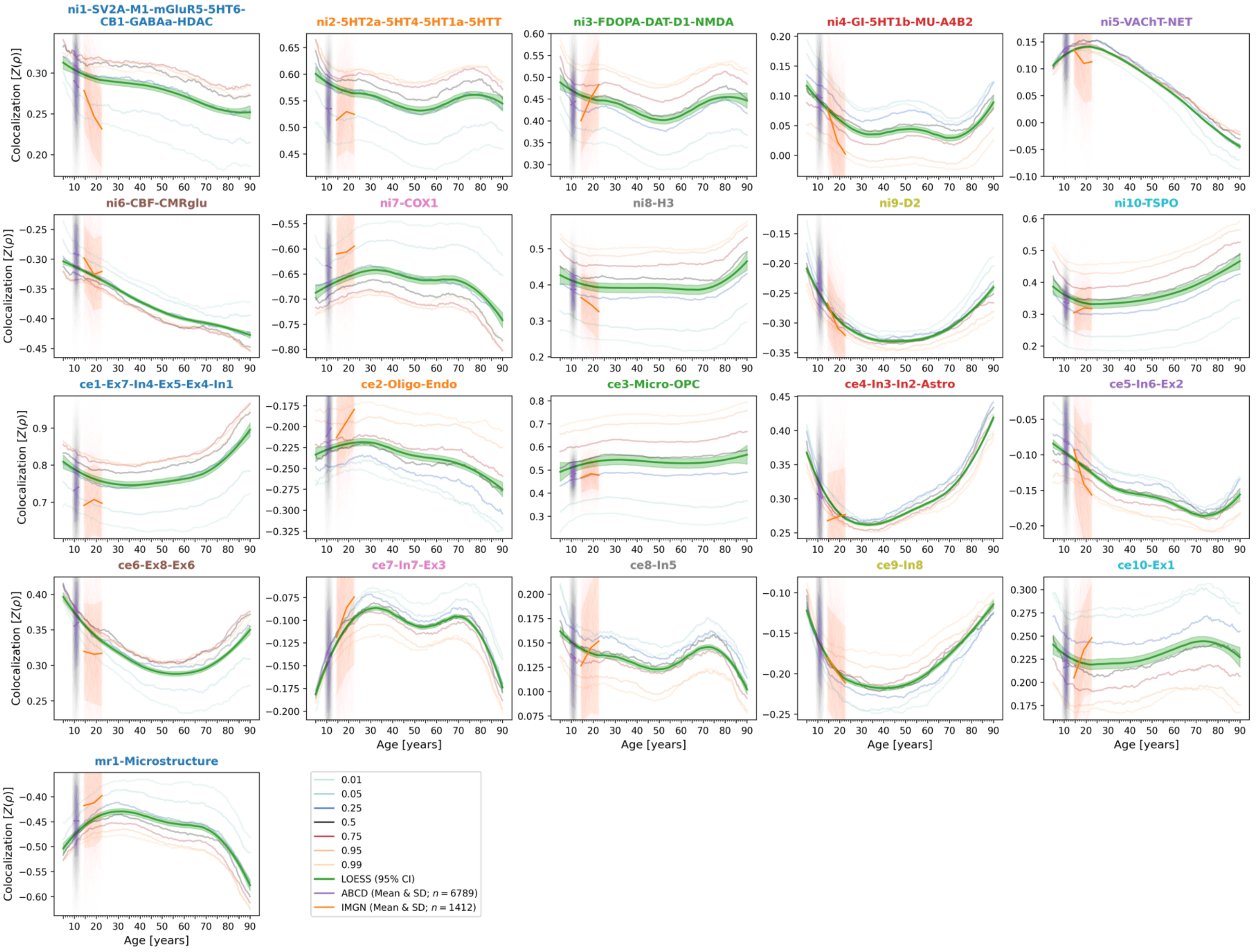

**Fig. S7: Spatial colocalization between modeled cross-sectional CT and neurobiological markers, including validation cohort data**

Lifespan trajectories of colocalization between neurobiological markers and modeled cross-sectional CT. Equivalent to Fig. 2 but with added individual subjects from ABCD and IMAGEN cohorts (**lines connecting individual timepoints**) with mean colocalization strength and standard deviations indicated by the **thick lines and shaded areas**. Source data are provided as a Source Data file.

#### A: Comparison of colocalization trajectories

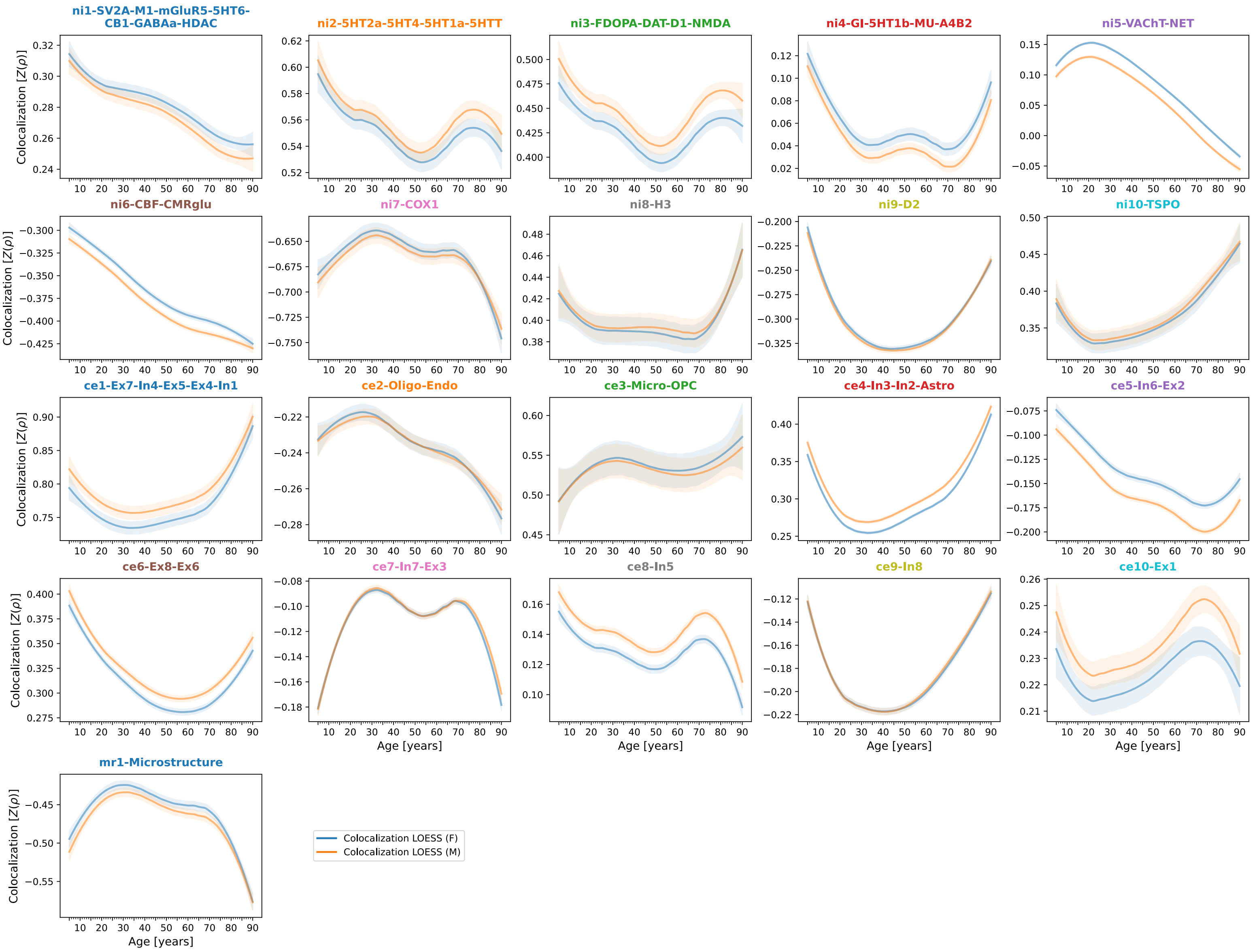

#### B: Female only

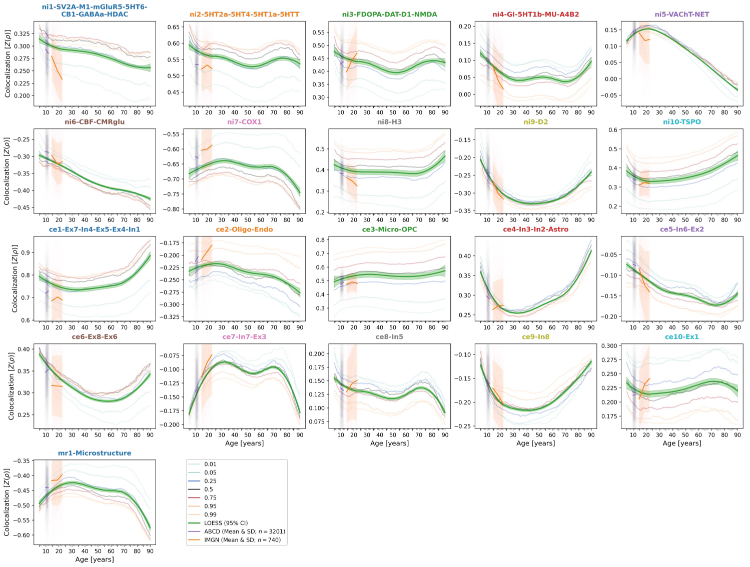

#### C: Male only

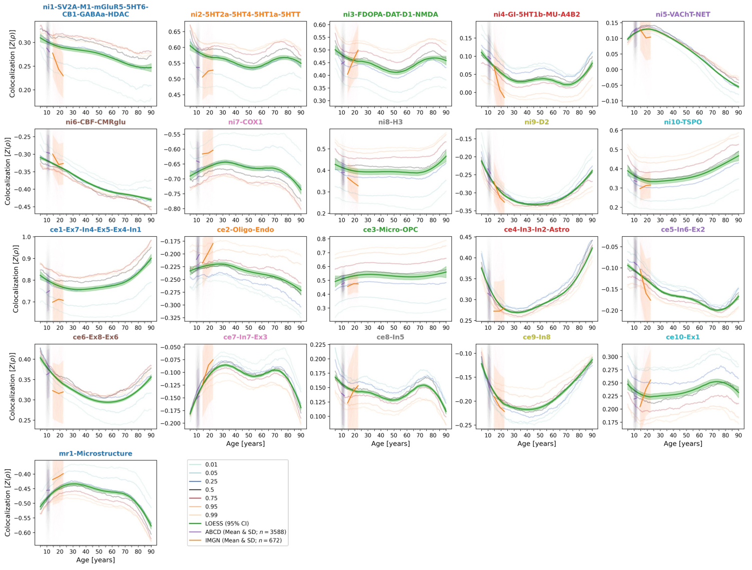

**Fig. S8: Spatial colocalization between cross-sectional modeled CT and neurobiological markers, compared between sexes**

Lifespan trajectories of colocalization between neurobiological markers and modeled cross-sectional CT. **(A)** Comparison of LOESS lines as in Figs. 2 and S7 but estimated on either female or male subjects only. **(B)** and **(C)**: Results for modeled CT and ABCD/IMAGEN data, separately by sex. Plot elements as in Fig. S7. **Abbreviations**: CT = cortical thickness, MRI = magnetic resonance imaging, LOESS = locally estimated scatterplot smoothing. Source data are provided as a Source Data file.

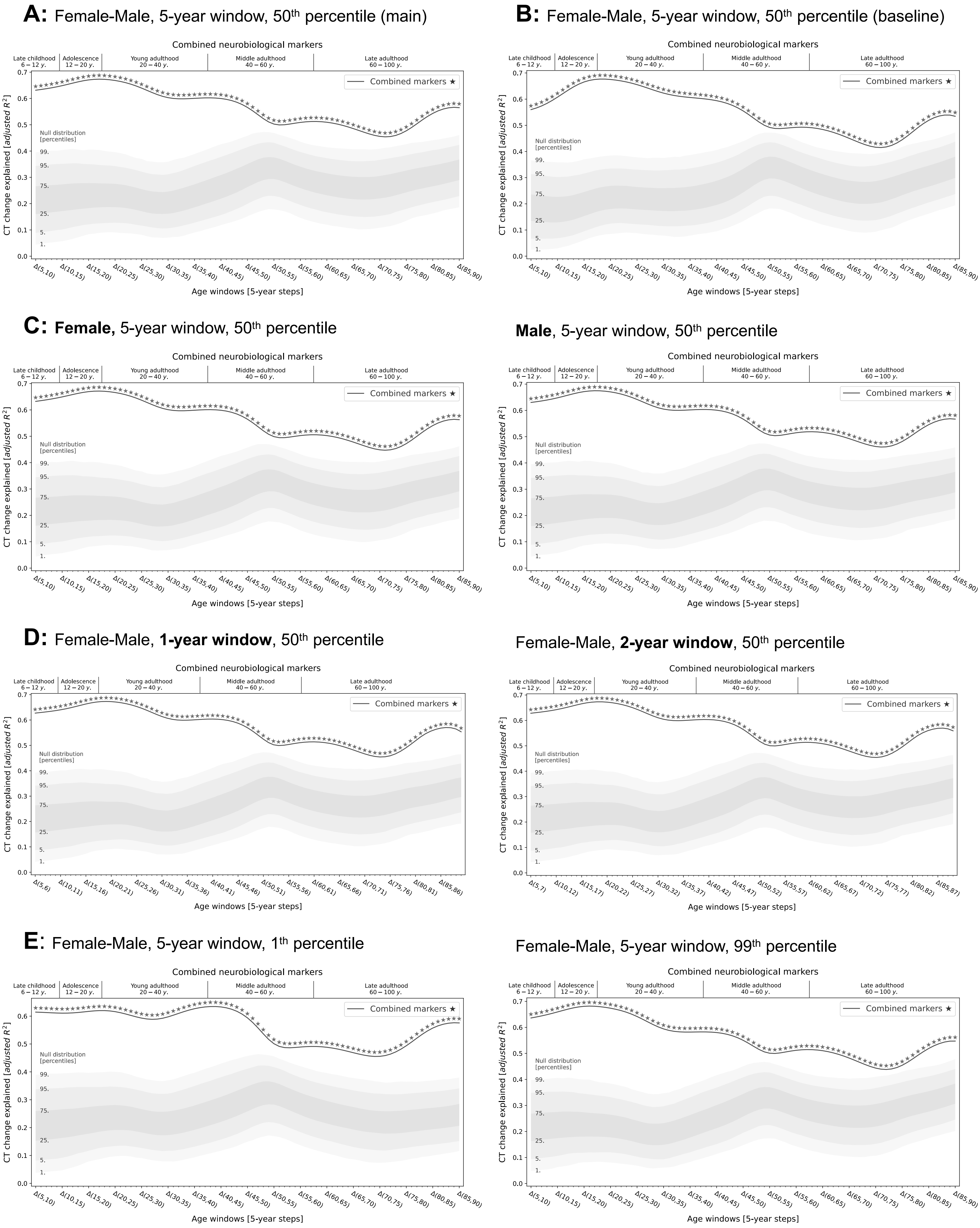

A: Female-Male, 5-year window, 50th percentile (main)
B: Female-Male, 5-year window, 50th percentile (baseline)
C: Female, 5-year window, 50th percentile
Male, 5-year window, 50th percentile
D: Female-Male, 1-year window, 50th percentile
Female-Male, 2-year window, 50th percentile
E: Female-Male, 5-year window, 1th percentile
Female-Male, 5-year window, 99th percentile
Combined neurobiological markers
Late childhood 6 − 12 y.
Adolescence 12 − 20 y.
Young adulthood 20 − 40 y.
Middle adulthood 40 − 60 y.
Late adulthood 60 − 100 y.
Combined markers ★
Null distribution [percentiles]
99.
95.
75.
25.
5.
1.
CT change explained [adjusted R²]
Age windows [5-year steps]

**Fig. S9: Joint multivariate regression result in different analysis cases**

---

**(A)** Multivariate regression on cortical thickness change across the lifespan using all predictors, neuroimaging and mRNA expression combined, settings as in main analyses. **(B)** Results after correcting the change data for baseline modeled CT. **(C)** Results for only male or only female modeled CT data. **(D)** Results with sliding window length of 1 or 2 years instead of 5 years. **(E)** Results for first or 99$^{th}$ instead of 50$^{th}$ percentile modeled CT data. Please refer to main Fig. 3 for descriptions of the panel elements. Source data are provided as a Source Data file.

---

#### Female-Male, 5-year window, 50th percentile **(main)**

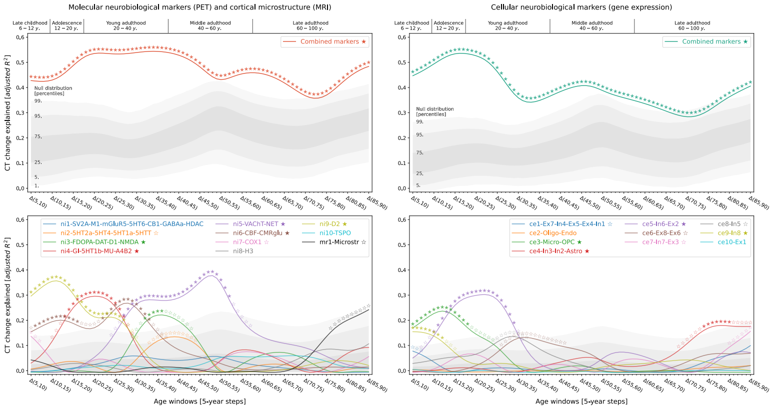

#### Female-Male, 5-year window, 50th percentile **(baseline)**

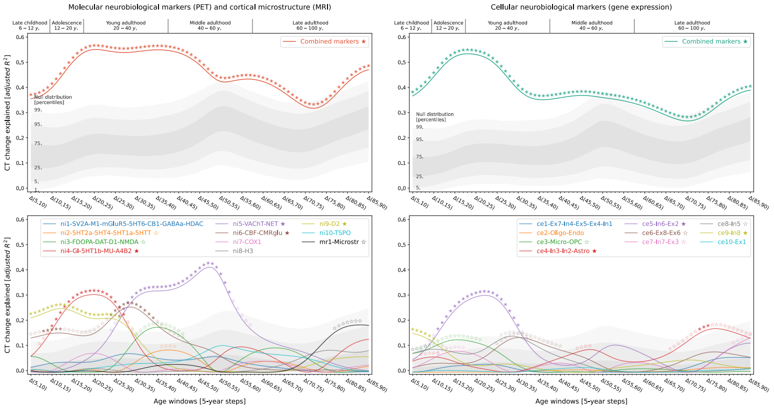

#### **Female**, 5-year window, 50th percentile

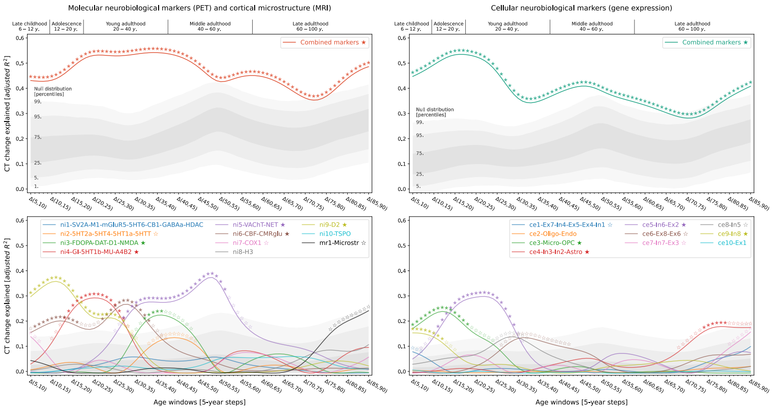

#### **Male**, 5-year window, 50th percentile

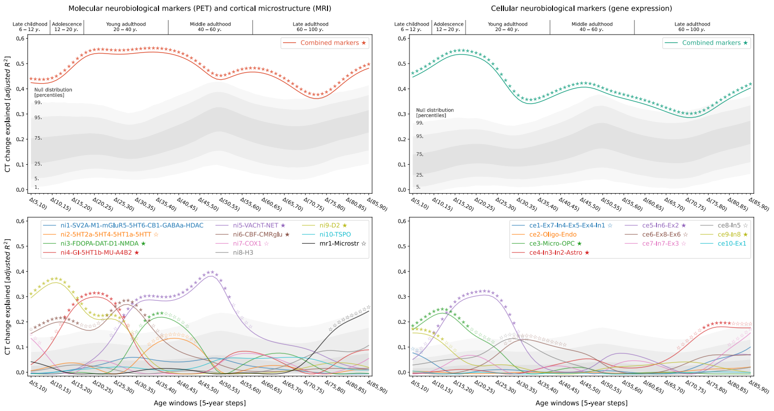

#### Female-Male, **1-year window**, 50th percentile

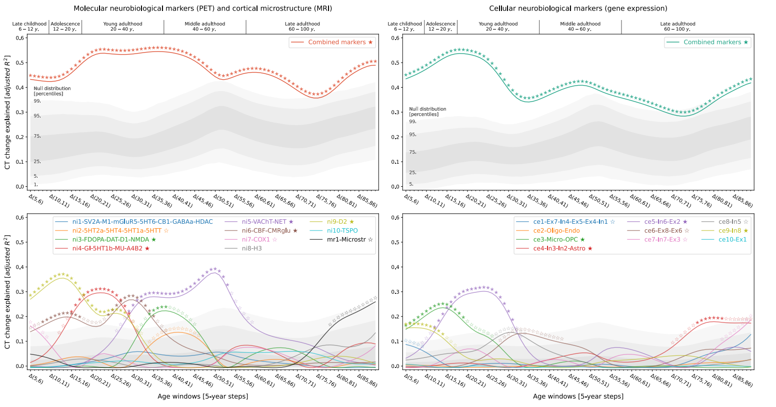

#### Female-Male, **2-year window**, 50th percentile

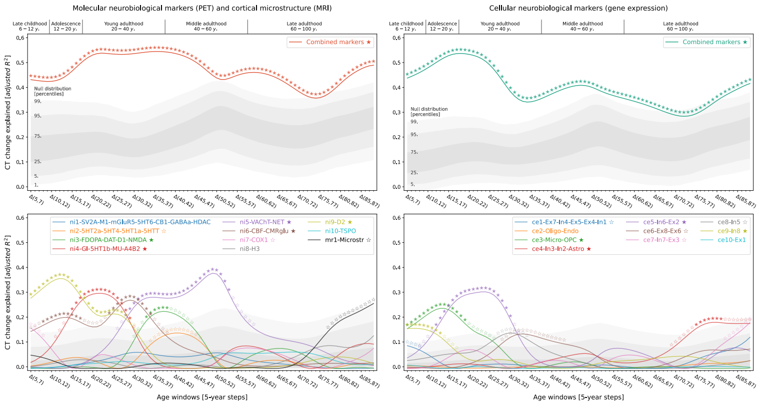

#### Female-Male, 5-year window, **1th percentile**

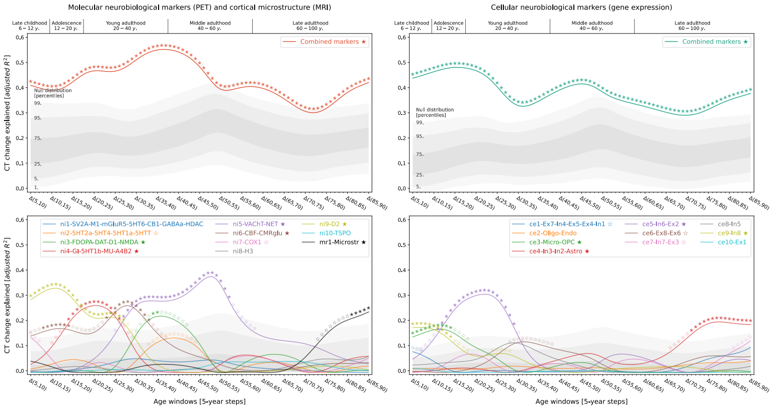

#### Female-Male, 5-year window, **99th percentile**

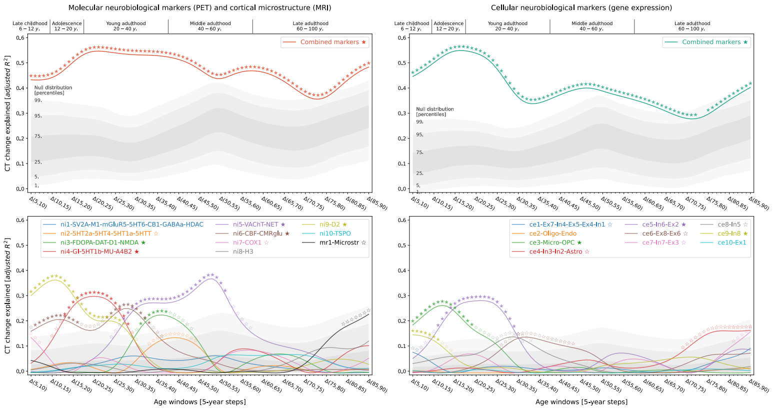

**Fig. S10: Influence of modeled sex, sliding window length, and cortical thickness reference percentile on main regression results**

Univariate and multivariate regression results for the cases outlined in Fig. S9. Please refer to main Fig. 3 for descriptions of the panel elements. Source data are provided as a Source Data file.

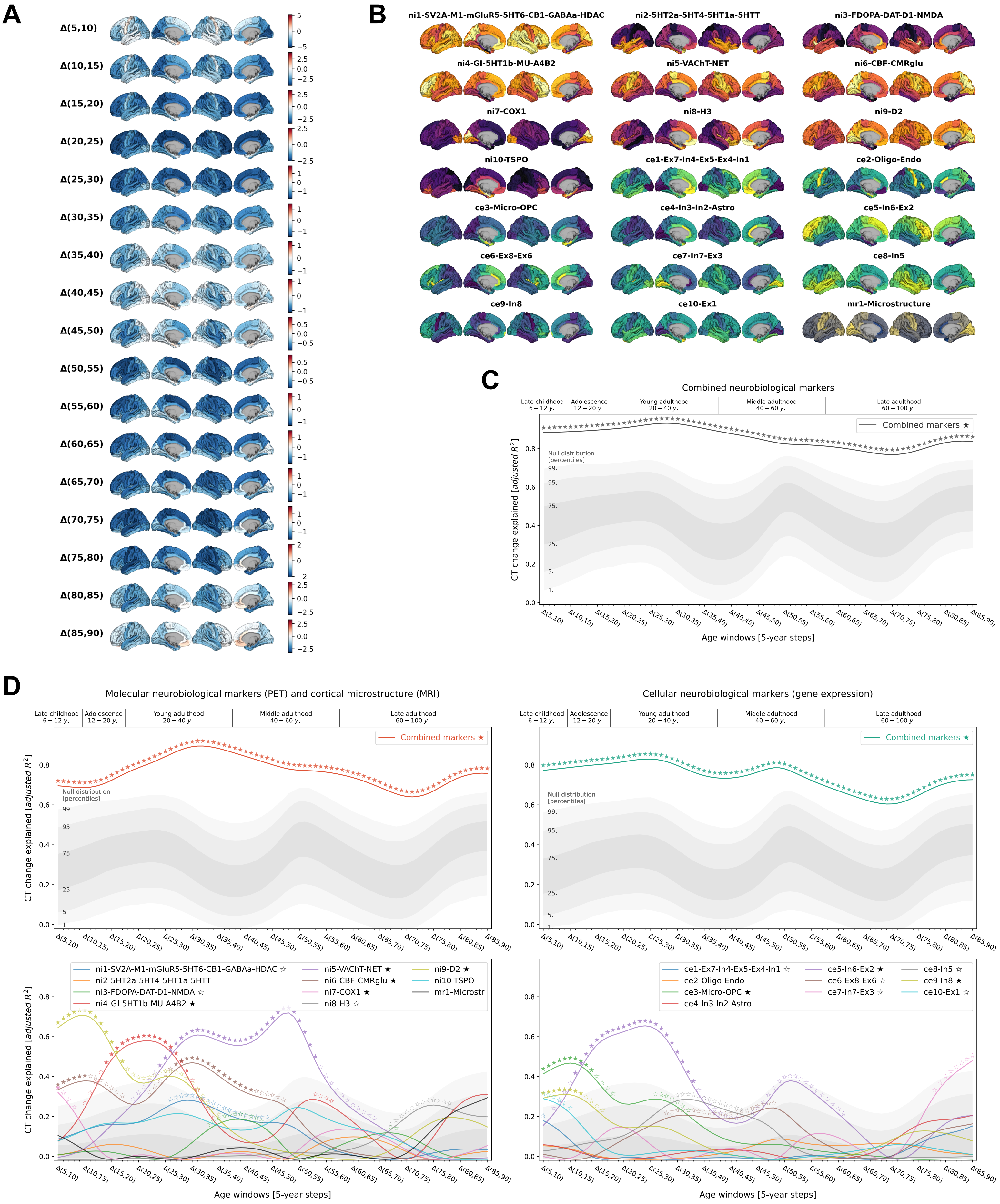
A
Δ(5,10)
Δ(10,15)
Δ(15,20)
Δ(20,25)
Δ(25,30)
Δ(30,35)
Δ(35,40)
Δ(40,45)
Δ(45,50)
Δ(50,55)
Δ(55,60)
Δ(60,65)
Δ(65,70)
Δ(70,75)
Δ(75,80)
Δ(80,85)
Δ(85,90)
B
ni1-SV2A-M1-mGluR5-5HT6-CB1-GABAa-HDAC
ni2-5HT2a-5HT4-5HT1a-5HTT
ni3-FDOPA-DAT-D1-NMDA
ni4-GI-5HT1b-MU-A4B2
ni5-VAChT-NET
ni6-CBF-CMRglu
ni7-COX1
ni8-H3
ni9-D2
ni10-TSPO
ce1-Ex7-In4-Ex5-Ex4-In1
ce2-Oligo-Endo
ce3-Micro-OPC
ce4-In3-In2-Astro
ce5-In6-Ex2
ce6-Ex8-Ex6
ce7-In7-Ex3
ce8-In5
ce9-In8
ce10-Ex1
mr1-Microstructure
C
Combined neurobiological markers
Late childhood 6 – 12 y.
Adolescence 12 – 20 y.
Young adulthood 20 – 40 y.
Middle adulthood 40 – 60 y.
Late adulthood 60 – 100 y.
Combined markers ★
Null distribution [percentiles]
CT change explained [adjusted R²]
Age windows [5-year steps]
D
Molecular neurobiological markers (PET) and cortical microstructure (MRI)
Cellular neurobiological markers (gene expression)
ni1-SV2A-M1-mGluR5-5HT6-CB1-GABAa-HDAC ☆
ni2-5HT2a-5HT4-5HT1a-5HTT
ni3-FDOPA-DAT-D1-NMDA ☆
ni4-GI-5HT1b-MU-A4B2 ★
ni5-VAChT-NET ★
ni6-CBF-CMRglu ★
ni7-COX1 ★
ni8-H3 ☆
ni9-D2 ★
ni10-TSPO
mr1-Microstr
ce1-Ex7-In4-Ex5-Ex4-In1 ☆
ce2-Oligo-Endo
ce3-Micro-OPC ★
ce4-In3-In2-Astro
ce5-In6-Ex2 ★
ce6-Ex8-Ex6 ☆
ce7-In7-Ex3 ☆
ce8-In5 ☆
ce9-In8 ★
ce10-Ex1 ☆
CT change explained [adjusted R²]
Age windows [5-year steps]

**Fig. S11: Replication of explained modeled CT change results using the Desikan-Killiany parcellation**

As the Desikan-Killiany parcellation[41] is the used more often in the literature, we transformed the Destrieux-parcellated data used in our study to the Desikan-Killiany parcels and repeated the first step of our regression analyses workflow. **(A)** and **(B)** show the Desikan-Killiany-transformed modeled CT change and factor-level neurobiological marker maps, respectively. **(C)** and **(D)** correspond to results shown in Figs. 3, 4, S9, and S10. Source data are provided as a Source Data file.

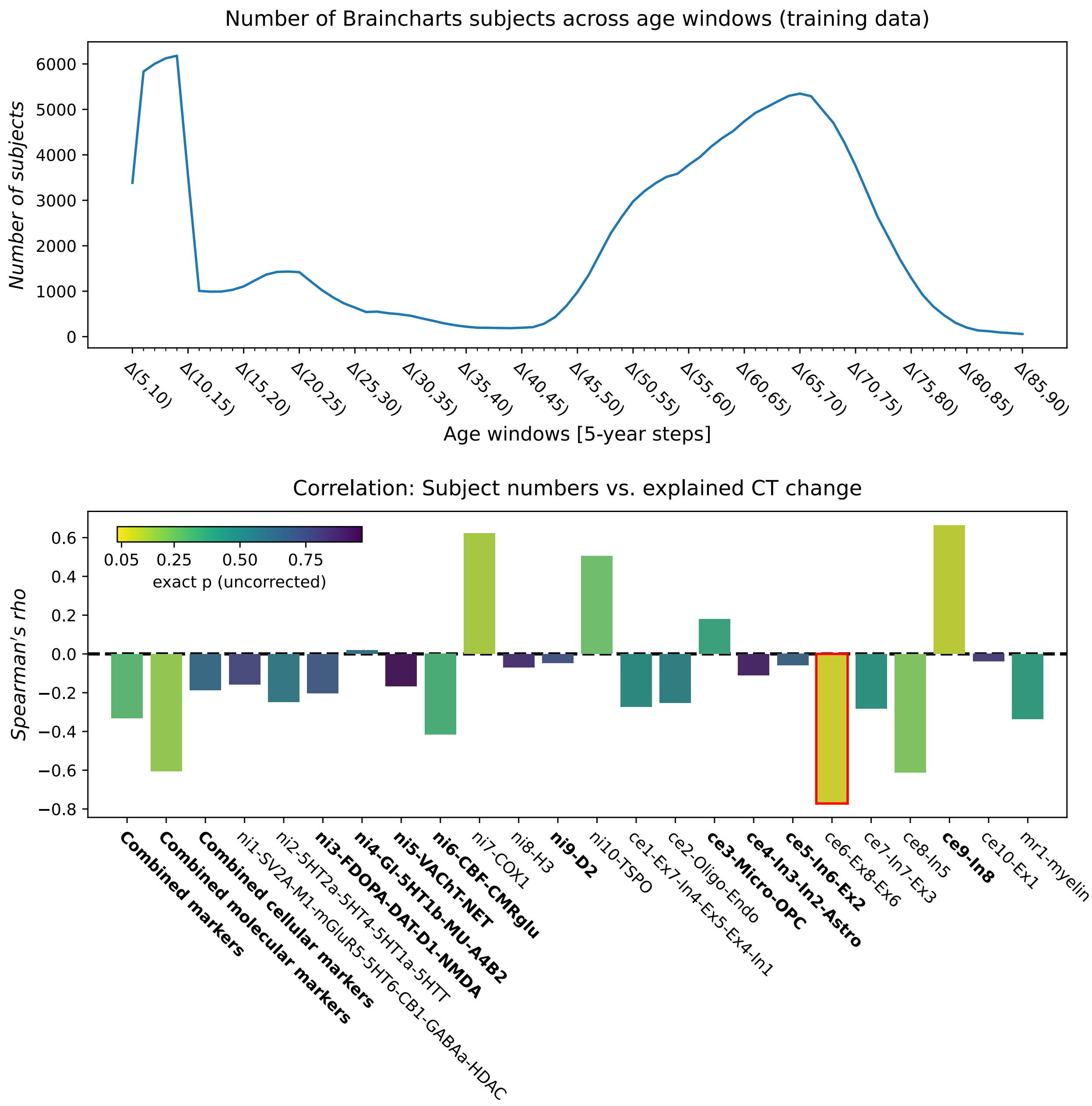

**Fig. S12: Potential influence of the non-uniform Braincharts age distribution on modeled CT change variance explained**

To test if the non-uniform age distribution of the Braincharts cohort could have influenced our results regarding the extent to which modeled CT change patterns were explained, we correlated the number of Braincharts subjects that fell into each studied age window (5-year windows, 1-year steps; **upper panel**) with the variance explained by CT change regression models across age windows. Spearman's rho estimates are shown in the **lower panel**, with **colors** marking the exact *p* value of each orrelation obtained from a correlation null distribution based on the same set of null maps from which the modeled CT change *p* values were estimated (n = 10,000). No CT change association with FDR-significant markers (c.f., Figs. 3 and 4; marked with **bold font**) was correlated to the age distribution. Source data are provided as a Source Data file.

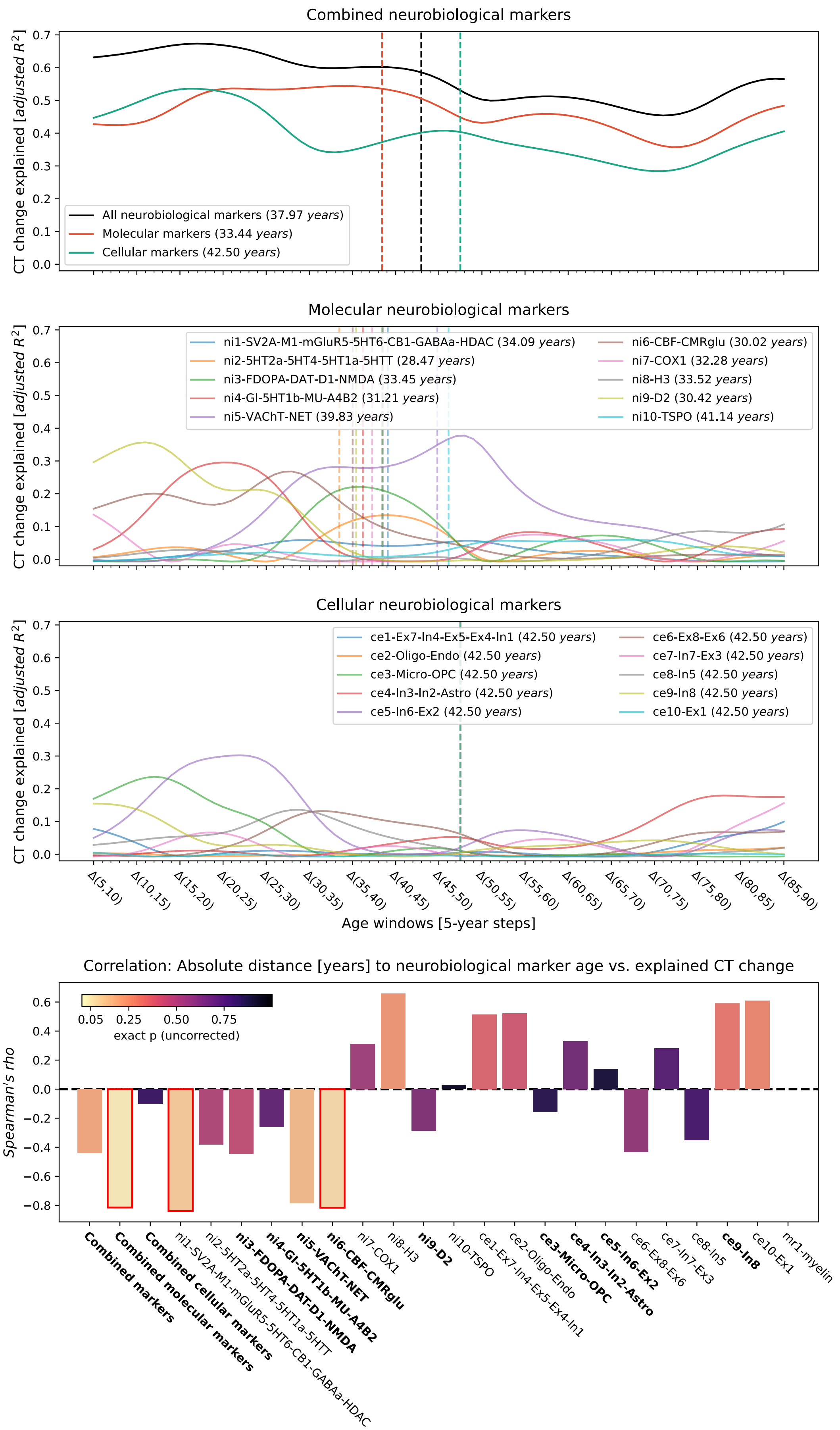
Combined neurobiological markers
CT change explained [adjusted R²]
All neurobiological markers (37.97 years)
Molecular markers (33.44 years)
Cellular markers (42.50 years)
Molecular neurobiological markers
ni1-SV2A-M1-mGluR5-5HT6-CB1-GABAa-HDAC (34.09 years)
ni2-5HT2a-5HT4-5HT1a-5HTT (28.47 years)
ni3-FDOPA-DAT-D1-NMDA (33.45 years)
ni4-GI-5HT1b-MU-A4B2 (31.21 years)
ni5-VAChT-NET (39.83 years)
ni6-CBF-CMRglu (30.02 years)
ni7-COX1 (32.28 years)
ni8-H3 (33.52 years)
ni9-D2 (30.42 years)
ni10-TSPO (41.14 years)
Cellular neurobiological markers
ce1-Ex7-In4-Ex5-Ex4-In1 (42.50 years)
ce2-Oligo-Endo (42.50 years)
ce3-Micro-OPC (42.50 years)
ce4-In3-In2-Astro (42.50 years)
ce5-In6-Ex2 (42.50 years)
ce6-Ex8-Ex6 (42.50 years)
ce7-In7-Ex3 (42.50 years)
ce8-In5 (42.50 years)
ce9-In8 (42.50 years)
ce10-Ex1 (42.50 years)
Δ(5,10) Δ(10,15) Δ(15,20) Δ(20,25) Δ(25,30) Δ(30,35) Δ(35,40) Δ(40,45) Δ(45,50) Δ(50,55) Δ(55,60) Δ(60,65) Δ(65,70) Δ(70,75) Δ(75,80) Δ(80,85) Δ(85,90)
Age windows [5-year steps]
Correlation: Absolute distance [years] to neurobiological marker age vs. explained CT change
Spearman's rho
exact p (uncorrected)
Combined markers
Combined molecular markers
Combined cellular markers
ni1-SV2A-M1-mGluR5-5HT6-CB1-GABAa-HDAC
ni2-5HT2a-5HT4-5HT1a-5HTT
ni3-FDOPA-DAT-D1-NMDA
ni4-GI-5HT1b-MU-A4B2
ni5-VAChT-NET
ni6-CBF-CMRglu
ni7-COX1
ni8-H3
ni9-D2
ni10-TSPO
ce1-Ex7-In4-Ex5-Ex4-In1
ce2-Oligo-Endo
ce3-Micro-OPC
ce4-In3-In2-Astro
ce5-In6-Ex2
ce6-Ex8-Ex6
ce7-In7-Ex3
ce8-In5
ce9-In8
ce10-Ex1
mr1-myelin

**Fig. S13: Potential influence of approximated neurobiological marker source age on modeled CT change variance explained**

To test if the neurobiological marker source ages could have influenced our results regarding the extent to which modeled CT change patterns were explained, we correlated the absolute distance of each marker's approximated age to each time window with the variance explained by modeled CT change regression models across age windows. Resulting Spearman's rho estimates are shown in the lower panel, with colors marking the exact p value of each orrelation obtained from a correlation null distribution based on the same set of null maps from which the modeled CT change p values were estimated (n = 10,000). We only found significant negative correlations and no systematic pattern, providing no evidence for concerning confounds. Source data are provided as a Source Data file.

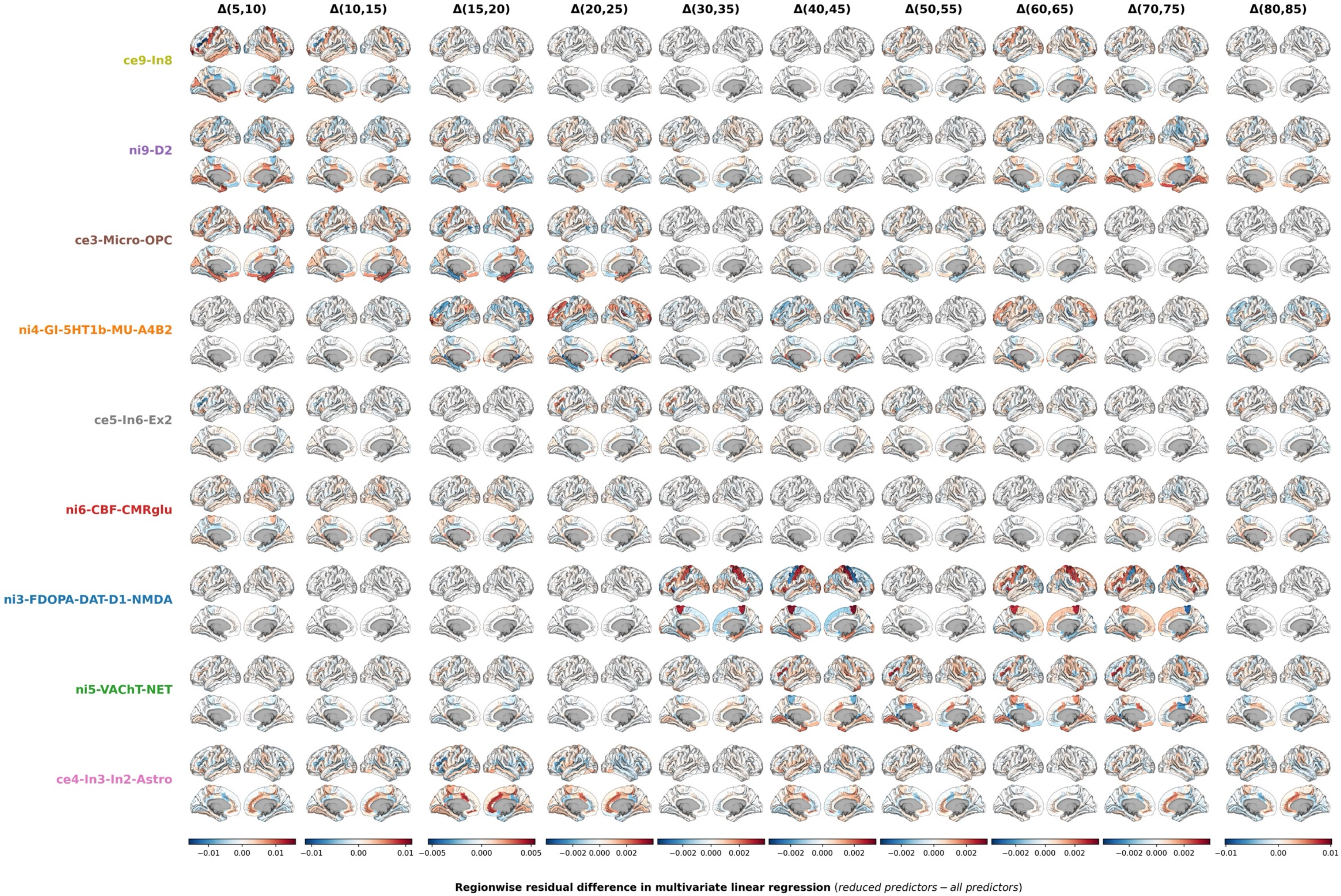

Δ(5,10)
Δ(10,15)
Δ(15,20)
Δ(20,25)
Δ(30,35)
Δ(40,45)
Δ(50,55)
Δ(60,65)
Δ(70,75)
Δ(80,85)
ce9-In8
ni9-D2
ce3-Micro-OPC
ni4-GI-5HT1b-MU-A4B2
ce5-In6-Ex2
ni6-CBF-CMRglu
ni3-FDOPA-DAT-D1-NMDA
ni5-VAChT-NET
ce4-In3-In2-Astro
−0.01 0.00 0.01
−0.01 0.00 0.01
−0.005 0.000 0.005
−0.002 0.000 0.002
−0.002 0.000 0.002
−0.002 0.000 0.002
−0.002 0.000 0.002
−0.002 0.000 0.002
−0.002 0.000 0.002
−0.01 0.00 0.01
Regionwise residual difference in multivariate linear regression (reduced predictors − all predictors)

**Fig. S14: Contribution of individual brain regions to explained lifespan cortical thickness change**

Parcellated brain plots show residual differences as estimated in the main dominance analyses. Residual differences were calculated for each predictor x as the difference between prediction errors resulting from a multivariate linear regression with all 7 predictors included and the prediction errors under exclusion of x. Source data are provided as a Source Data file.

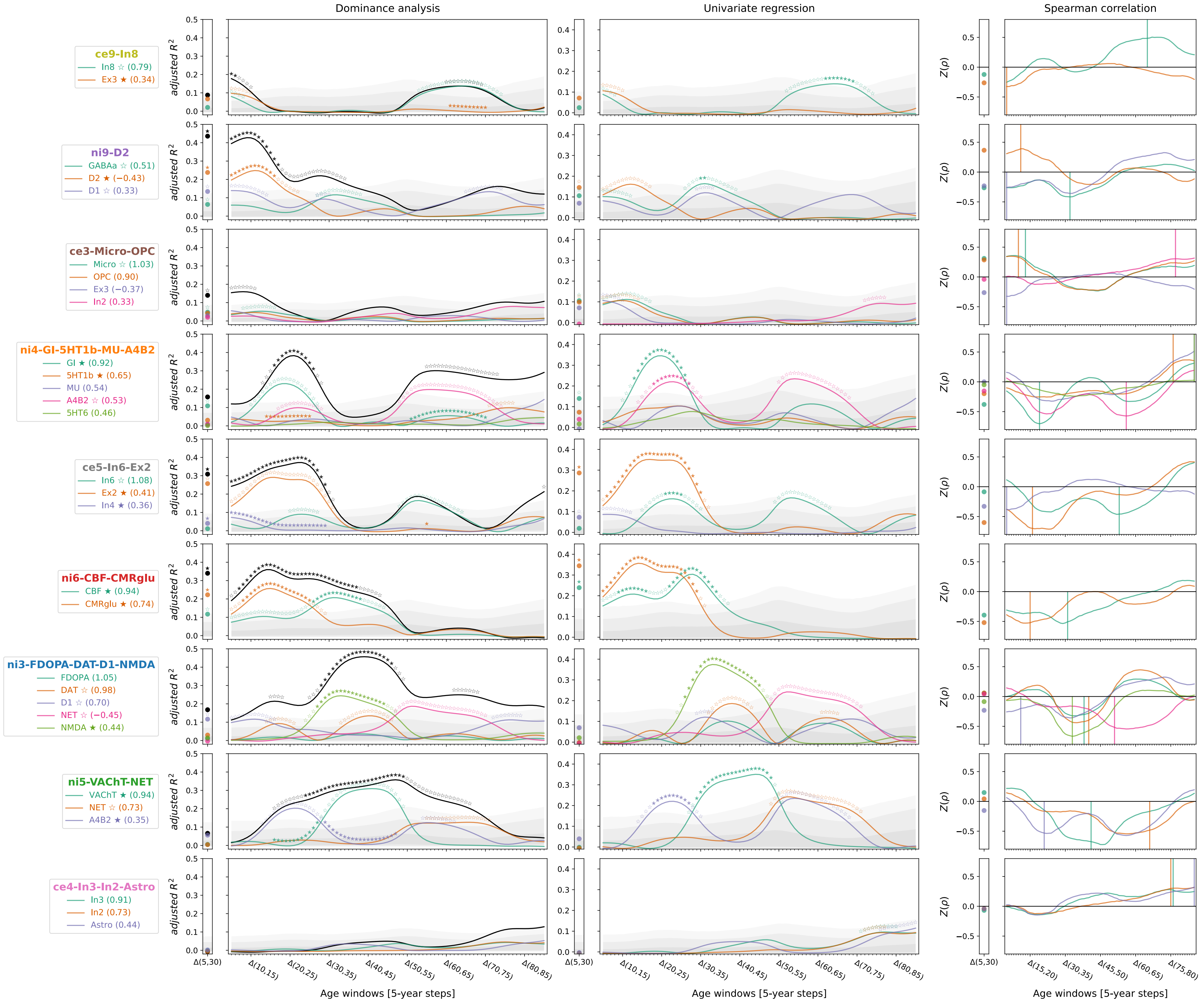
Dominance analysis
Univariate regression
Spearman correlation
ce9-In8
In8 ☆ (0.79)
Ex3 ★ (0.34)
ni9-D2
GABAa ☆ (0.51)
D2 ★ (−0.43)
D1 ☆ (0.33)
ce3-Micro-OPC
Micro ☆ (1.03)
OPC (0.90)
Ex3 (−0.37)
In2 (0.33)
ni4-GI-5HT1b-MU-A4B2
GI ★ (0.92)
5HT1b ★ (0.65)
MU (0.54)
A4B2 ☆ (0.53)
5HT6 (0.46)
ce5-In6-Ex2
In6 ☆ (1.08)
Ex2 ★ (0.41)
In4 ★ (0.36)
ni6-CBF-CMRglu
CBF ★ (0.94)
CMRglu ★ (0.74)
ni3-FDOPA-DAT-D1-NMDA
FDOPA (1.05)
DAT ☆ (0.98)
D1 ☆ (0.70)
NET ☆ (−0.45)
NMDA ★ (0.44)
ni5-VAChT-NET
VAChT ★ (0.94)
NET ☆ (0.73)
A4B2 ★ (0.35)
ce4-In3-In2-Astro
In3 (0.91)
In2 (0.73)
Astro (0.44)
adjusted $R^2$
$Z(\rho)$
Δ(5,30)
Δ(10,15)
Δ(20,25)
Δ(30,35)
Δ(40,45)
Δ(50,55)
Δ(60,65)
Δ(70,75)
Δ(80,85)
Δ(15,20)
Δ(45,50)
Δ(75,80)
Age windows [5-year steps]

**Fig. S15: Individual dominance analyses using original neurobiological atlases**

To determine if the factor-level predictors appropriately captured the original multimodal atlases, sets of spatial association analyses were calculated, predicting modeled CT change across the lifespan from the original maps most closely associated to each factor. For each factor, the 5 original atlases that loaded most strongly on the factor were selected if their loading exceeded a threshold of .3. The **first column** shows the result of stepwise dominance analyses (**black line** = combined $R^2$), the **second column** shows independent single linear regressions, and the **third column** depicts the colocalization pattern between modeled CT change and original predictors to illustrate the sign of the spatial association. **Gray shades** in the first two columns show the null distributions associated with the individual factor-level $R^2$ values. Source data are provided as a Source Data file.

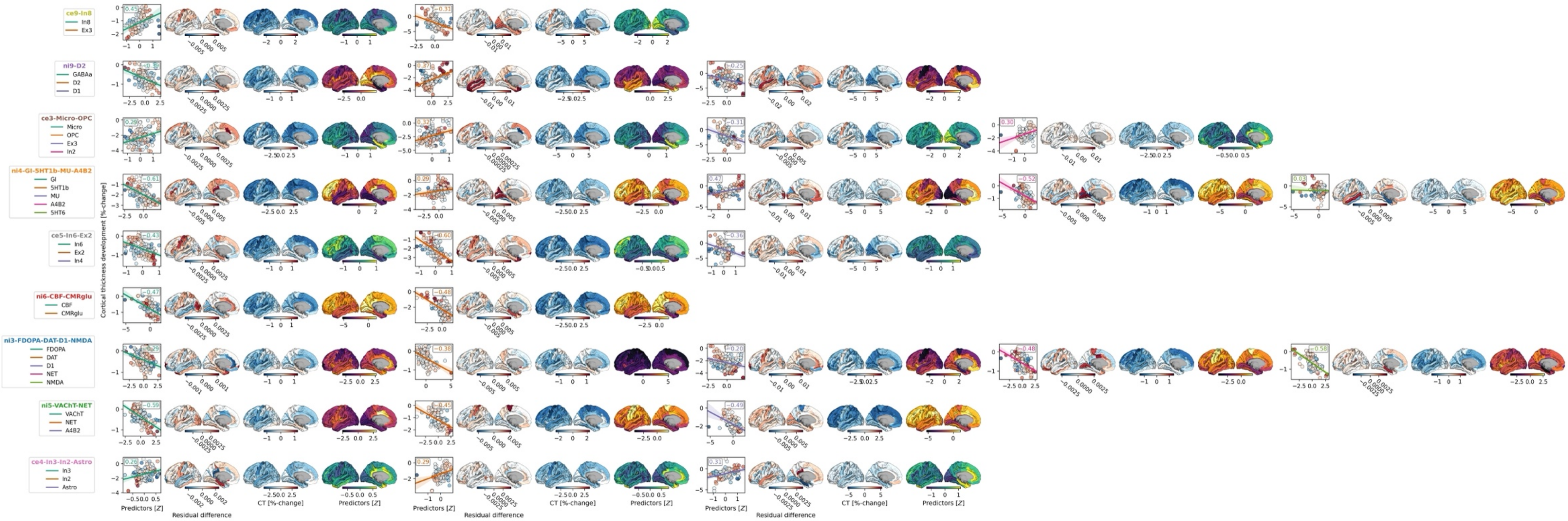

**Fig. S16: Cortical distributions of residual differences, cortical thickness changes, and average values of the original atlases associated to each selected predictor**

This figure corresponds to each row of Fig. S15 and mirrors the layout of Fig. 6. Please see these for descriptions of the panel elements. Source data are provided as a Source Data file.

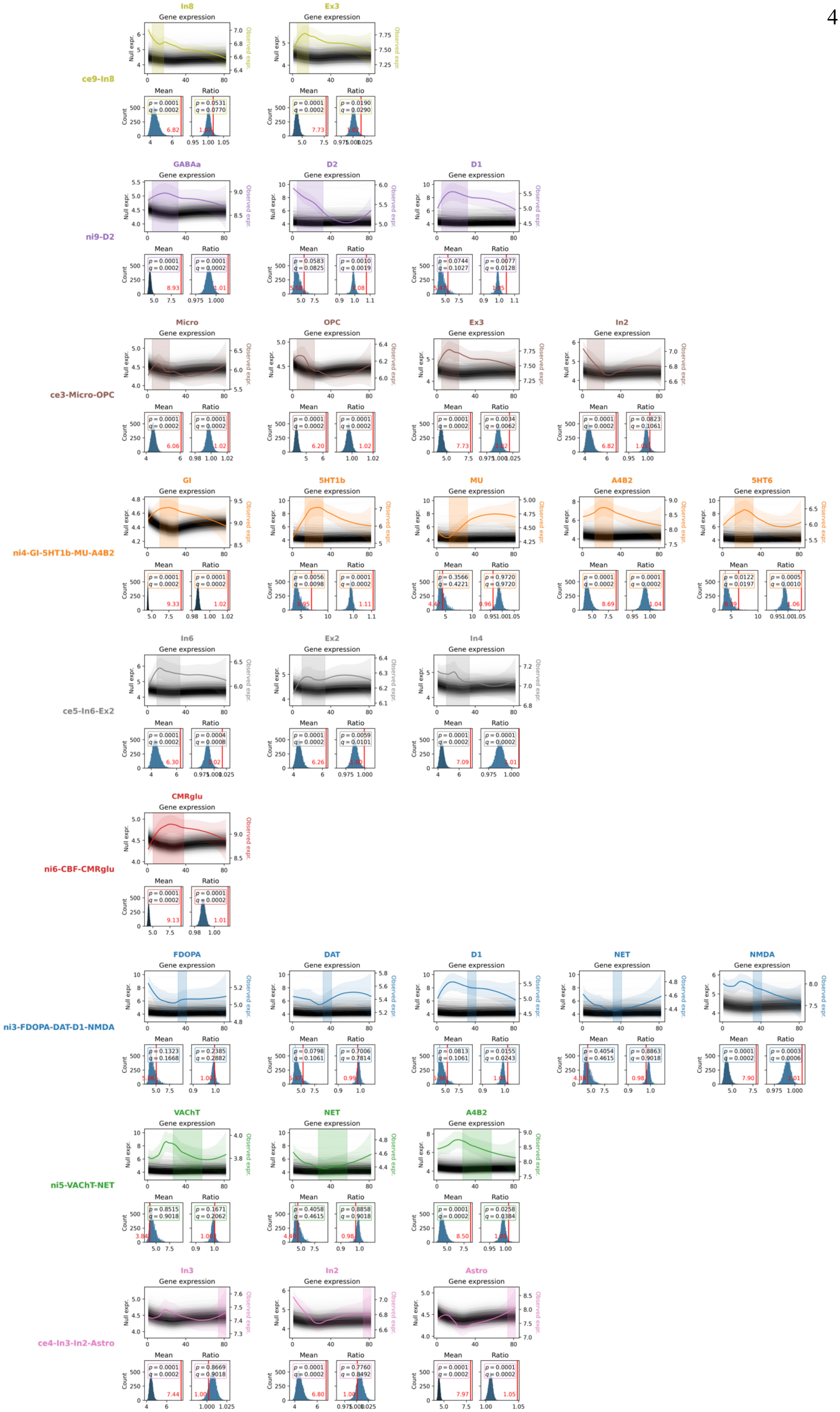
ce9-In8
In8
Ex3
Gene expression
Null expr.
Observed expr.
Mean
Ratio
Count
p = 0.0001 q = 0.0002
p = 0.0531 q = 0.0770
p = 0.0001 q = 0.0002
p = 0.0190 q = 0.0290
ni9-D2
GABAa
D2
D1
p = 0.0001 q = 0.0002
p = 0.0001 q = 0.0002
p = 0.0583 q = 0.0825
p = 0.0010 q = 0.0019
p = 0.0744 q = 0.1027
p = 0.0077 q = 0.0128
ce3-Micro-OPC
Micro
OPC
Ex3
In2
p = 0.0001 q = 0.0002
p = 0.0001 q = 0.0002
p = 0.0001 q = 0.0002
p = 0.0001 q = 0.0002
p = 0.0001 q = 0.0002
p = 0.0034 q = 0.0062
p = 0.0001 q = 0.0002
p = 0.0823 q = 0.1061
ni4-GI-5HT1b-MU-A4B2
GI
5HT1b
MU
A4B2
5HT6
p = 0.0001 q = 0.0002
p = 0.0001 q = 0.0002
p = 0.0056 q = 0.0098
p = 0.0001 q = 0.0002
p = 0.3566 q = 0.4221
p = 0.9720 q = 0.9720
p = 0.0001 q = 0.0002
p = 0.0001 q = 0.0002
p = 0.0122 q = 0.0197
p = 0.0005 q = 0.0010
ce5-In6-Ex2
In6
Ex2
In4
p = 0.0001 q = 0.0002
p = 0.0004 q = 0.0008
p = 0.0001 q = 0.0002
p = 0.0059 q = 0.0101
p = 0.0001 q = 0.0002
p = 0.0001 q = 0.0002
ni6-CBF-CMRglu
CMRglu
p = 0.0001 q = 0.0002
p = 0.0001 q = 0.0002
ni3-FDOPA-DAT-D1-NMDA
FDOPA
DAT
D1
NET
NMDA
p = 0.1323 q = 0.1668
p = 0.2385 q = 0.2882
p = 0.0798 q = 0.1061
p = 0.7006 q = 0.7814
p = 0.0813 q = 0.1061
p = 0.0155 q = 0.0243
p = 0.4054 q = 0.4615
p = 0.8863 q = 0.9018
p = 0.0001 q = 0.0002
p = 0.0003 q = 0.0006
ni5-VAChT-NET
VAChT
NET
A4B2
p = 0.8515 q = 0.9018
p = 0.1671 q = 0.2062
p = 0.4058 q = 0.4615
p = 0.8858 q = 0.9018
p = 0.0001 q = 0.0002
p = 0.0258 q = 0.0384
ce4-In3-In2-Astro
In3
In2
Astro
p = 0.0001 q = 0.0002
p = 0.8669 q = 0.9018
p = 0.0001 q = 0.0002
p = 0.7760 q = 0.8492
p = 0.0001 q = 0.0002
p = 0.0001 q = 0.0002

**Fig. S17: Null trajectories and test null distributions for developmental gene expression data**

Detailed test results corresponding to main Fig. 7. For each neurobiological marker **(rows)** and each gene/ gene set associated to the original brain atlases **(columns per row)**, we show the observed gene expression trajectory **(colored, left y axis)**, and $n$ = 10,000 null trajectories **(grey, right y axis)**. The histograms below show null distribution for the mean expression during the significant CT period **(left)** and the ratio of mean expression during vs. outside the significant modeled CT period **(right)**. The **red lines and numbers** indicate observed mean/ ratio values; $p$ and $q$ indicate nominal and FDR-corrected empirical $p$ values. Source data are provided as a Source Data file.

#### A: ABCD

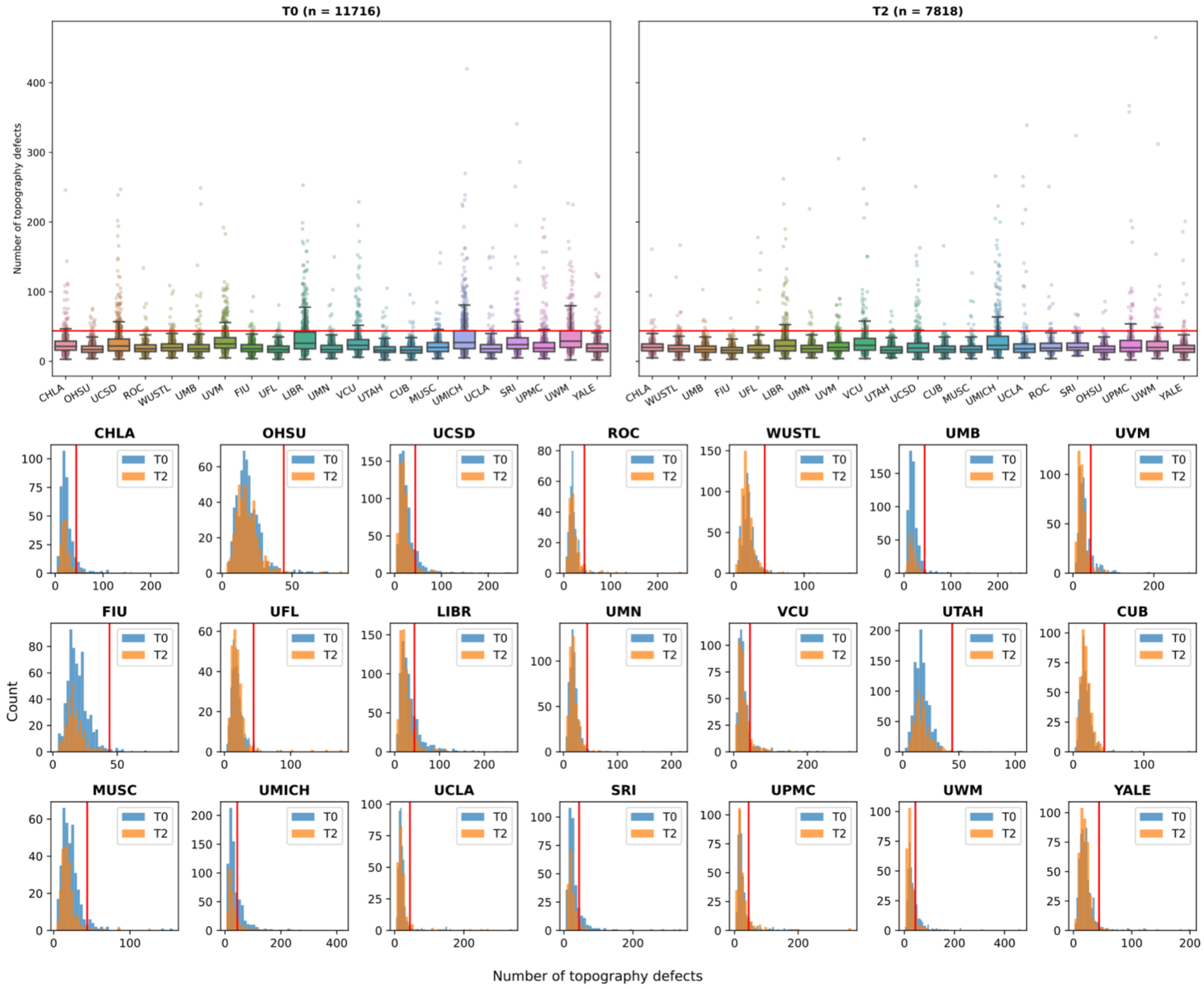

#### B: IMAGEN

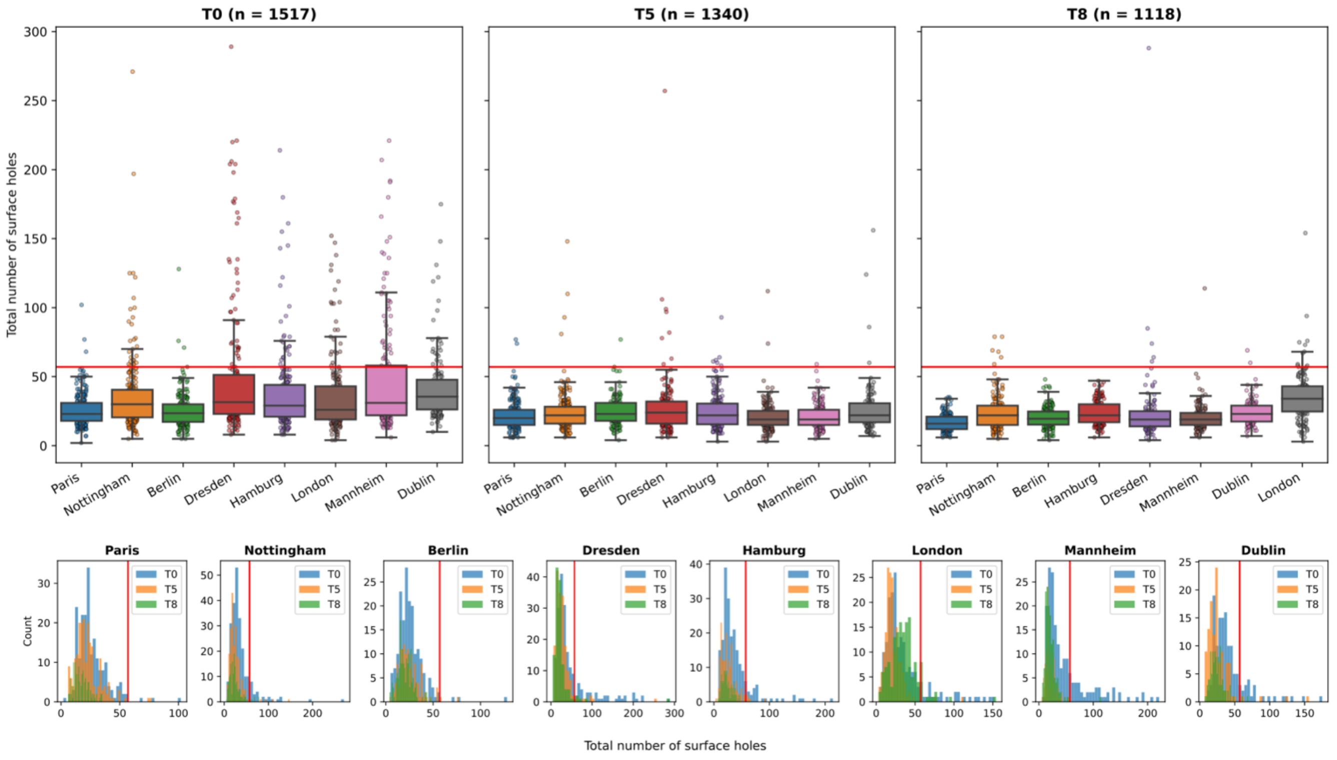

**Fig. S18: Euler number-based quality control of ABCD and IMAGEN data**

Distributions of Euler number-related metrics in ABCD **(A)** and IMAGEN **(B)** datasets at different study time points. The upper and lower sub-panels of A and B visualize the distributions of the quality control metrics in different ways. For IMAGEN, the sum of the two hemispheric Euler numbers, and for ABCD, the variable "apqc_smri_topo_ndefect" was used. Subjects were excluded if they exceeded a threshold of Q3 + 1.5 * IQR calculated across study time points within each study. Group-level source data are provided as a Source Data file.

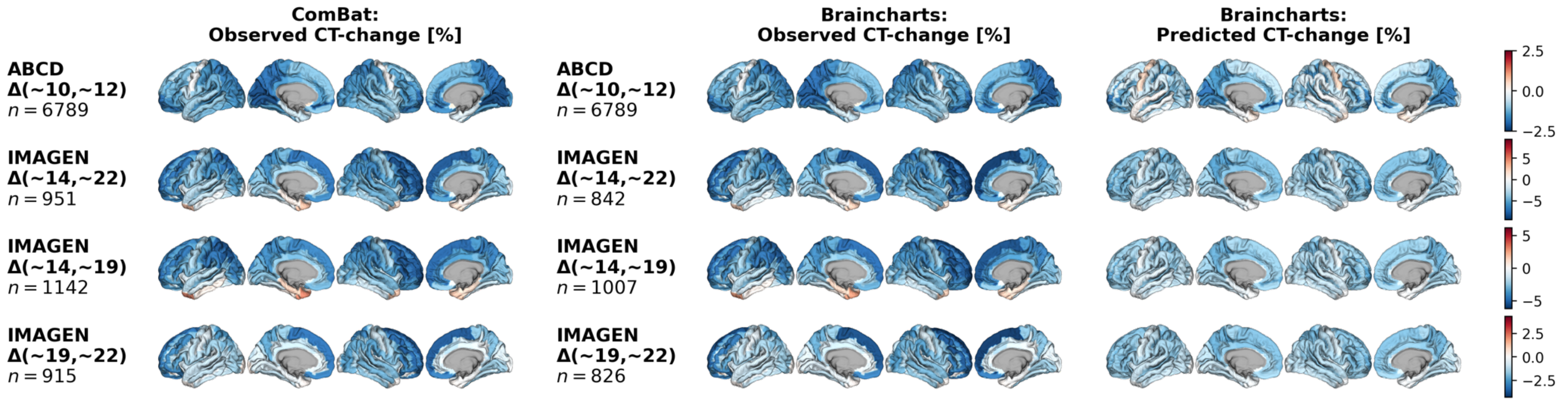

**Fig. S19: Observed and predicted ABCD and IMAGEN cohort-average cortical thickness change**

Average relative CT change in percent across four time periods in the ABCD and IMAGEN datasets as observed, after ComBat harmonization (**left**), as observed after adaptation to the Braincharts model (**middle**), and as predicted by the Braincharts model based on subject sex and age **(right)**. Sample sizes vary for IMAGEN subjects between ComBat-harmonized and Braincharts-adjusted/predicted data as some subjects were used for model adaptation and then dropped from the analyses. This is not the case for ABCD subjects as, here, solely baseline-only subjects were used for model adaptation. Source data are provided as a Source Data file.

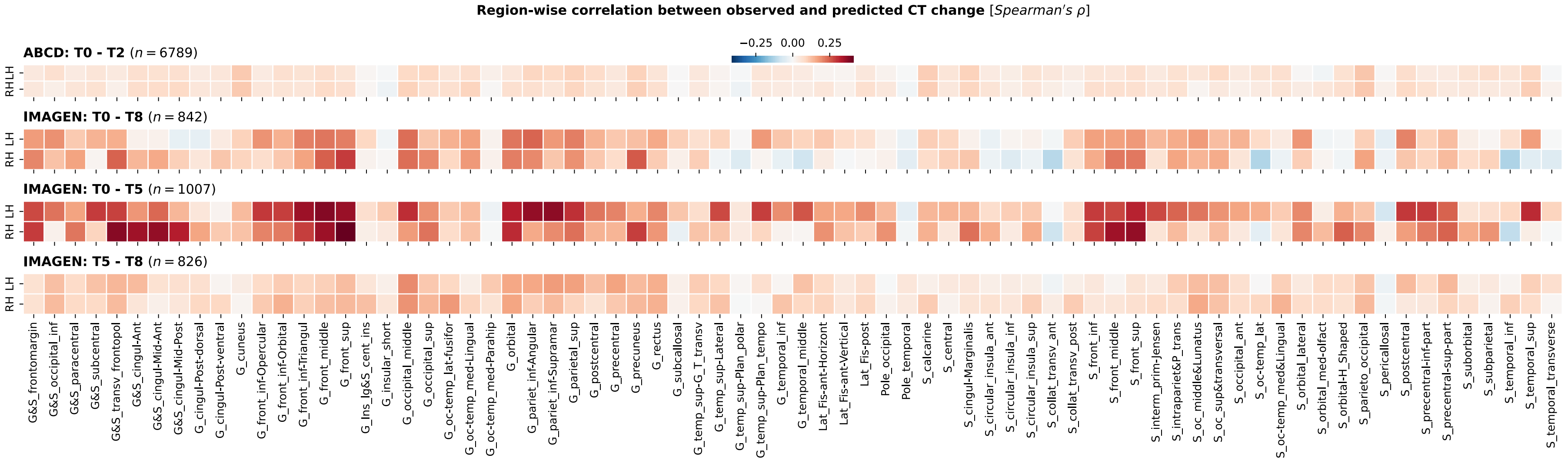

#### Fig. S20: Region-wise correlation between predicted and observed cortical thickness change

Regional correlation (Spearman's rho) between relative CT changes as predicted by the Braincharts model (percent-change of predicted CT values from timepoint one to timepoint two) and relative CT change as observed in ABCD and IMAGEN datasets across each time period, after adaptation to the Braincharts model. Predicted CT change was calculated for each subject individually based on their age and sex. Source data are provided as a Source Data file.

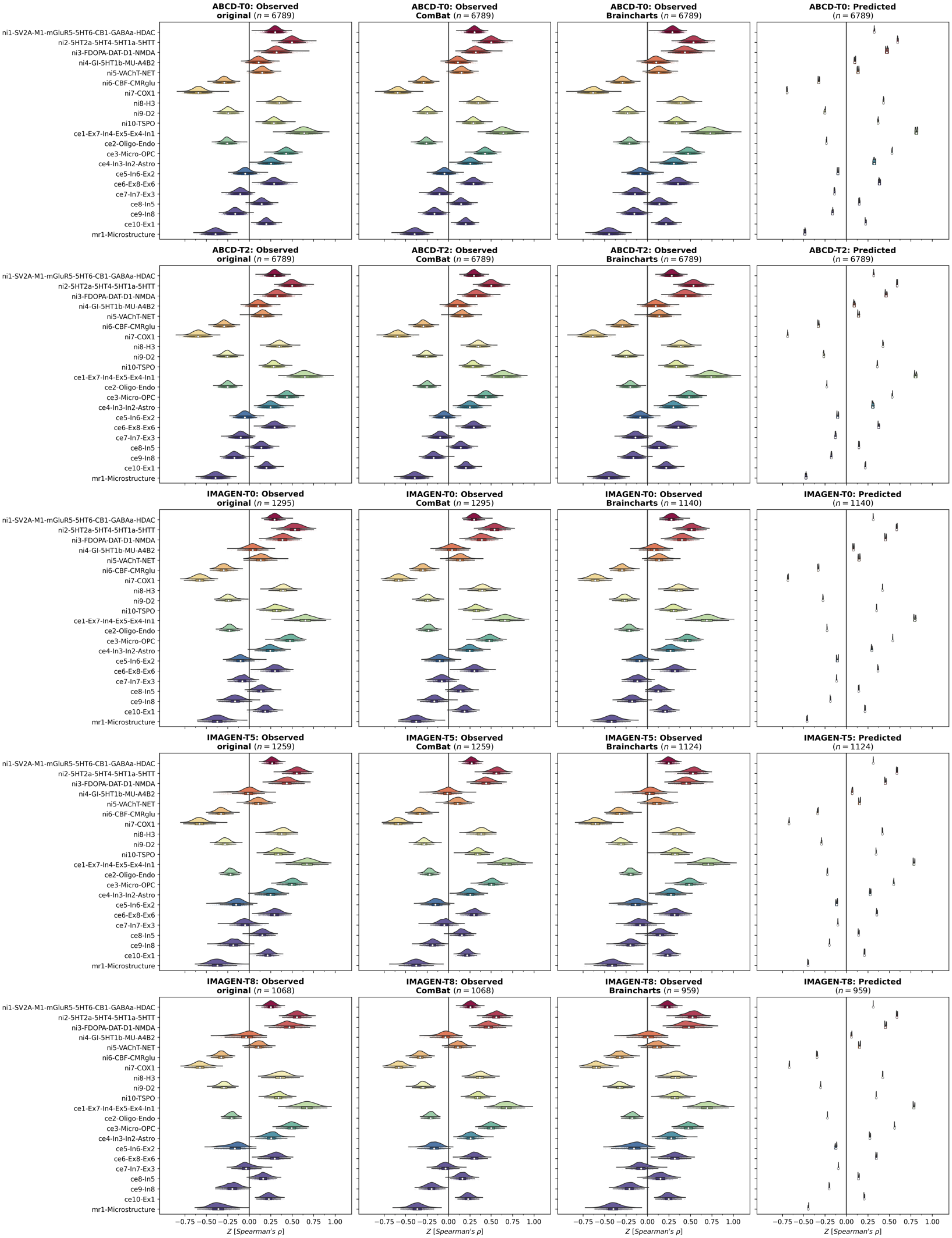
ABCD-T0: Observed original (n = 6789)
ABCD-T0: Observed ComBat (n = 6789)
ABCD-T0: Observed Brainchart (n = 6789)
ABCD-T0: Predicted (n = 6789)
ABCD-T2: Observed original (n = 6789)
ABCD-T2: Observed ComBat (n = 6789)
ABCD-T2: Observed Brainchart (n = 6789)
ABCD-T2: Predicted (n = 6789)
IMAGEN-T0: Observed original (n = 1295)
IMAGEN-T0: Observed ComBat (n = 1295)
IMAGEN-T0: Observed Brainchart (n = 1140)
IMAGEN-T0: Predicted (n = 1140)
IMAGEN-T5: Observed original (n = 1259)
IMAGEN-T5: Observed ComBat (n = 1259)
IMAGEN-T5: Observed Brainchart (n = 1124)
IMAGEN-T5: Predicted (n = 1124)
IMAGEN-T8: Observed original (n = 1068)
IMAGEN-T8: Observed ComBat (n = 1068)
IMAGEN-T8: Observed Brainchart (n = 959)
IMAGEN-T8: Predicted (n = 959)
ni1-SV2A-M1-mGluR5-5HT6-CB1-GABAa-HDAC
ni2-5HT2a-5HT4-5HT1a-5HTT
ni3-FDOPA-DAT-D1-NMDA
ni4-GI-5HT1b-MU-A4B2
ni5-VAChT-NET
ni6-CBF-CMRglu
ni7-COX1
ni8-H3
ni9-D2
ni10-TSPO
ce1-Ex7-In4-Ex5-Ex4-In1
ce2-Oligo-Endo
ce3-Micro-OPC
ce4-In3-In2-Astro
ce5-In6-Ex2
ce6-Ex8-Ex6
ce7-In7-Ex3
ce8-In5
ce9-In8
ce10-Ex1
mr1-Microstructure
−0.75
−0.50
−0.25
0.00
0.25
0.50
0.75
1.00
Z [Spearman's ρ]

**Fig. S21: Spatial colocalization between cross-sectional ABCD and IMAGEN cortical thickness data and multimodal neurobiological markers**

Spatial colocalization with multimodal predictors quantified as Z-transformed Spearman correlation for each ABCD and IMAGEN subject at each study time point (**rows**) based on the original CT data without harmonization (**first column**), observed CT data after ComBat-harmonization (**second column**), observed CT data after site-correction using the Braincharts model (**third column**, note the smaller sample sizes for the IMAGEN data), and CT data as predicted by the Braincharts model (**fourth column**). See also the trajectory plots (Figs. S7 and S8). Group-level source data are provided as a Source Data file.

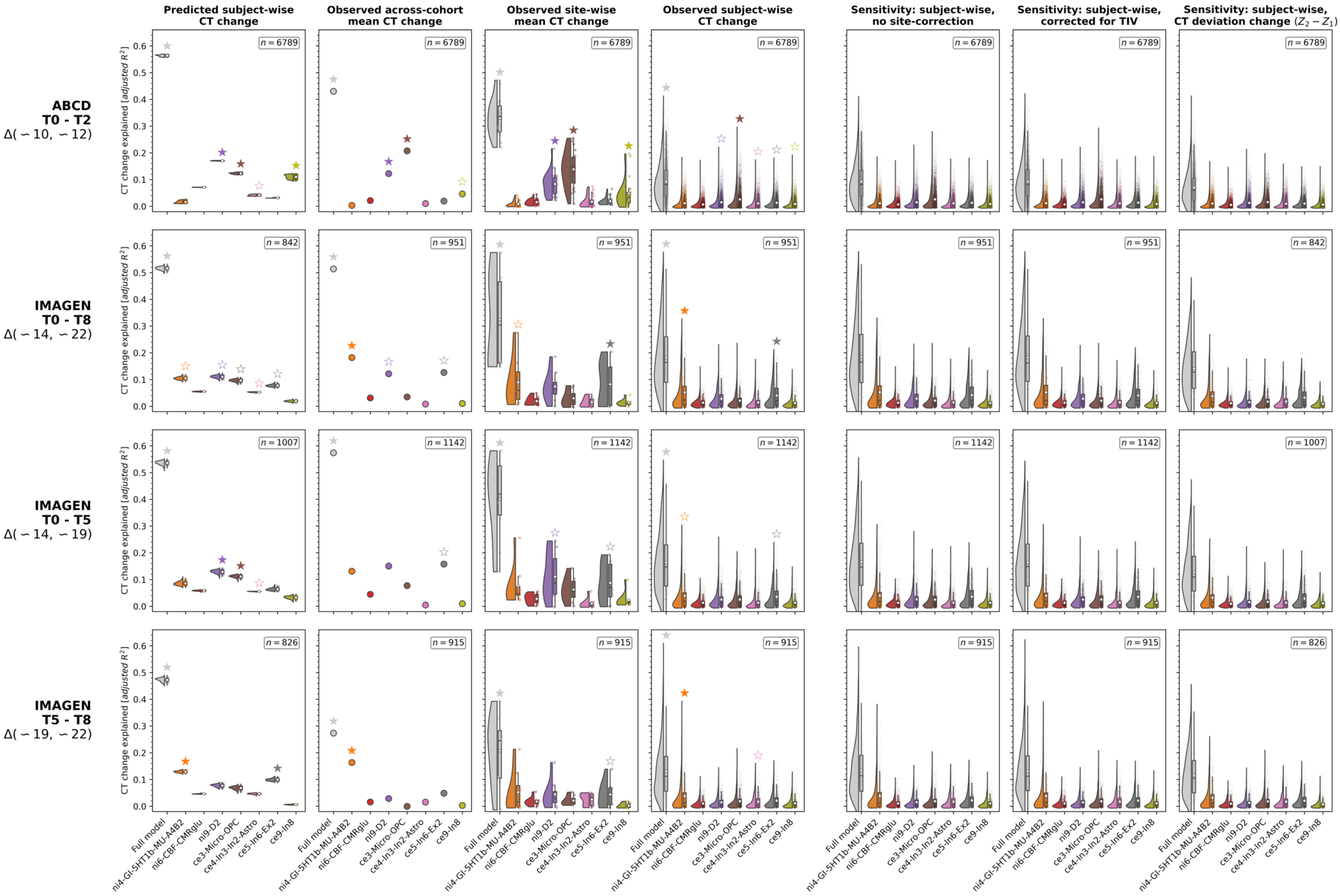
Predicted subject-wise CT change
Observed across-cohort mean CT change
Observed site-wise mean CT change
Observed subject-wise CT change
Sensitivity: subject-wise, no site-correction
Sensitivity: subject-wise, corrected for TIV
Sensitivity: subject-wise, CT deviation change ($Z_2 - Z_1$)
ABCD
T0 - T2
Δ(∼10, ∼12)
IMAGEN
T0 - T8
Δ(∼14, ∼22)
IMAGEN
T0 - T5
Δ(∼14, ∼19)
IMAGEN
T5 - T8
Δ(∼19, ∼22)
CT change explained [adjusted $R^2$]
$n = 6789$
$n = 842$
$n = 951$
$n = 1007$
$n = 1142$
$n = 826$
$n = 915$
Full model
ni4-GI-5HT1b-MU-A4B2
ni6-CBF-CMRglu
ni9-D2
ce3-Micro-OPC
ce4-In3-In2-Astro
ce5-In6-Ex2
ce9-In8

**Fig. S22: Detailed results of ABCD and IMAGEN validation analyses**

Explained CT change in ABCD and IMAGEN cohorts across 4 study time periods (**rows**) in different analysis settings. **Raincloud** plots and **scatters** show distributions of $R^2$ values resulting from cohort- or subject-wise dominance analyses. The **leftmost gray elements** depict the full model explained variance, the **right-sided colored elements** show the predictor-wise estimated *total dominance* statistics. For each subject- or cohort-wise analysis, the sum of predictor-wise $R^2$ values equals the full model value. **Stars** indicate statistical significance determined based on null distributions of $R^2$ values as estimated by rerunning regression analyses with predictor null maps (FDR-correction within each analysis/panel). See Fig. S23 for equivalent plots showing Spearman correlations. **(Column 1)** prediction result based on CT change as predicted by the Braincharts model. **(Column 2)** average CT change across each cohort, data after ComBat harmonization. **(Column 3)** average CT change across each site within each cohort, data after ComBar harmonization. **(Column 4)** CT change in single subjects, data after ComBat harmonization. **(Column 5, 6, and 7)** sensitivity analyses on subject-level data; **(5)** original values without site-correction to estimate effects of overfitting, **(6)** data after total intracranial volume was regressed from timepoint-wise estimates after ComBat harmonization, **(7)** change between deviation Z scores as estimated by the normative model. Group-level source data are provided as a Source Data file.

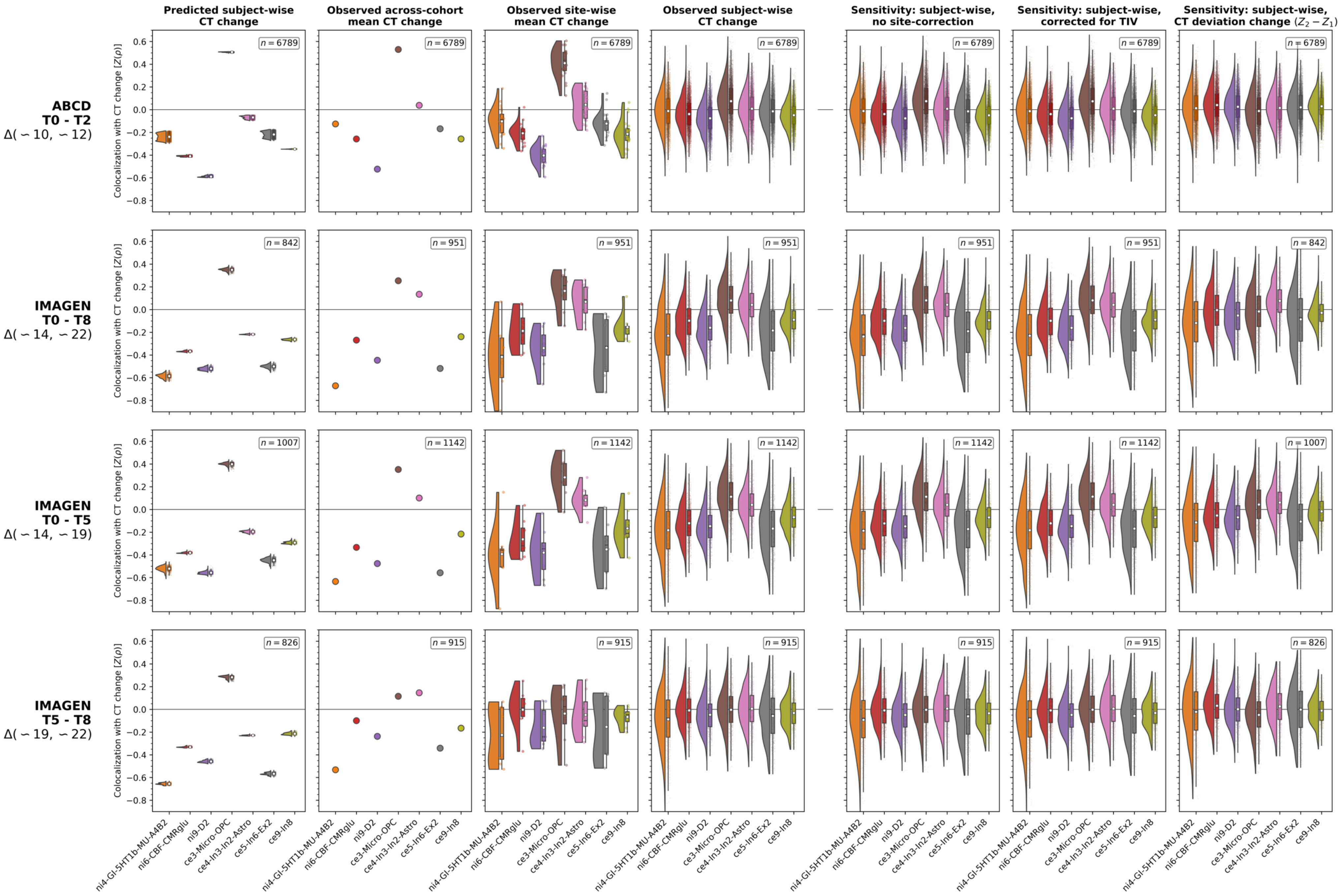

Predicted subject-wise CT change
Observed across-cohort mean CT change
Observed site-wise mean CT change
Observed subject-wise CT change
Sensitivity: subject-wise, no site-correction
Sensitivity: subject-wise, corrected for TIV
Sensitivity: subject-wise, CT deviation change ($Z_2 - Z_1$)
ABCD T0 - T2 Δ(∼10, ∼12)
IMAGEN T0 - T8 Δ(∼14, ∼22)
IMAGEN T0 - T5 Δ(∼14, ∼19)
IMAGEN T5 - T8 Δ(∼19, ∼22)
Colocalization with CT change [Z(ρ)]
n = 6789
n = 842
n = 951
n = 1007
n = 1142
n = 826
n = 915
ni4-GI-5HT1b-MU-A4B2
ni6-CBF-CMRglu
ni9-D2
ce3-Micro-OPC
ce4-In3-In2-Astro
ce5-In6-Ex2
ce9-In8

**Fig. S23: Detailed Spearman correlation results showing colocalization patterns between ABCD and IMAGEN CT change and multimodal predictors**

Add-on to Fig. S22. Instead of multivariate regression analyses, Spearman correlations were calculated, to capture the sign of the spatial associations. As of this demonstrational purpose, no significance tests were performed. Group-level source data are provided as a Source Data file.

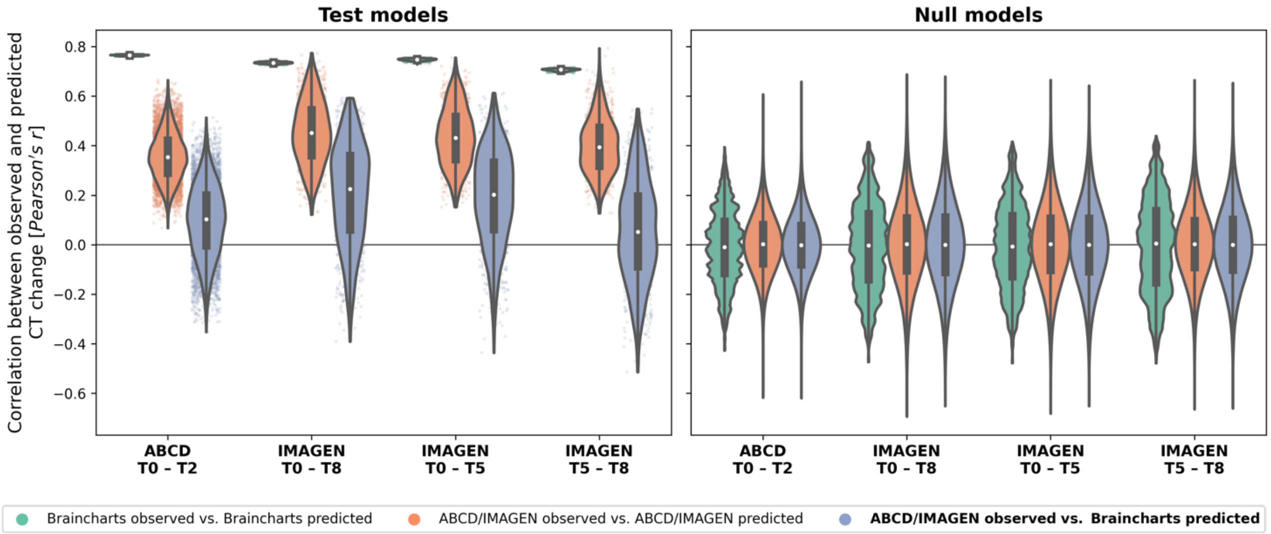

**Fig. S24: Generalization of CT change prediction models trained on normative single-subject data to observed data**

Evaluation of the generalizability of normative CT change prediction (= **blue violins**). The **y axis** shows regression model fit as the Pearson correlation between observed and predicted responses (i.e., CT change patterns). The **left panel** shows analyses based on the actual neurobiological marker brain maps, the **right panel** shows results based on permuted maps (1,000 iterations). **Green violins**: Fit of regression models estimated on each subject's *normative* CT change patterns as predicted by the Braincharts model. **Orange violins**: Fit of regression models estimated on each subject's observed CT change patterns. **Blue violins**: Fit of models estimated on the *normative* CT change patterns but applied to the *observed* CT change. The result indicates how well the normative model, as the "population average", performs in representing each subject's CT associations to neurobiological markers. Group-level source data are provided as a Source Data file.

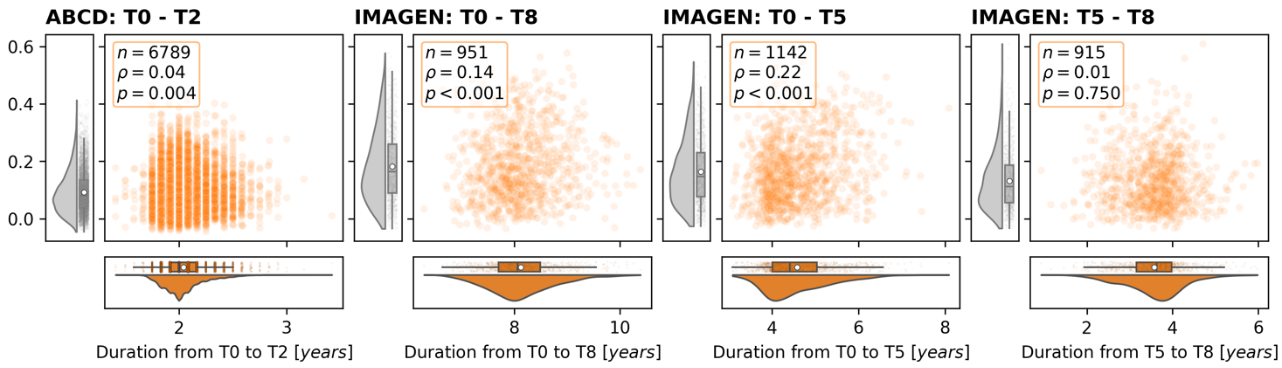

**Fig. S25: Effects of follow-up duration on total explained cortical thickness change**

Correlations between follow-up duration for each dataset and at each time period (**x axes**) with total explained CT change variance (**y axes**); full datasets after ComBat harmonization. **Boxes** in the upper corners of each panel contain Spearman's rho and the associated parametric *p* value for each bivariate association. Group-level source data are provided as a Source Data file.

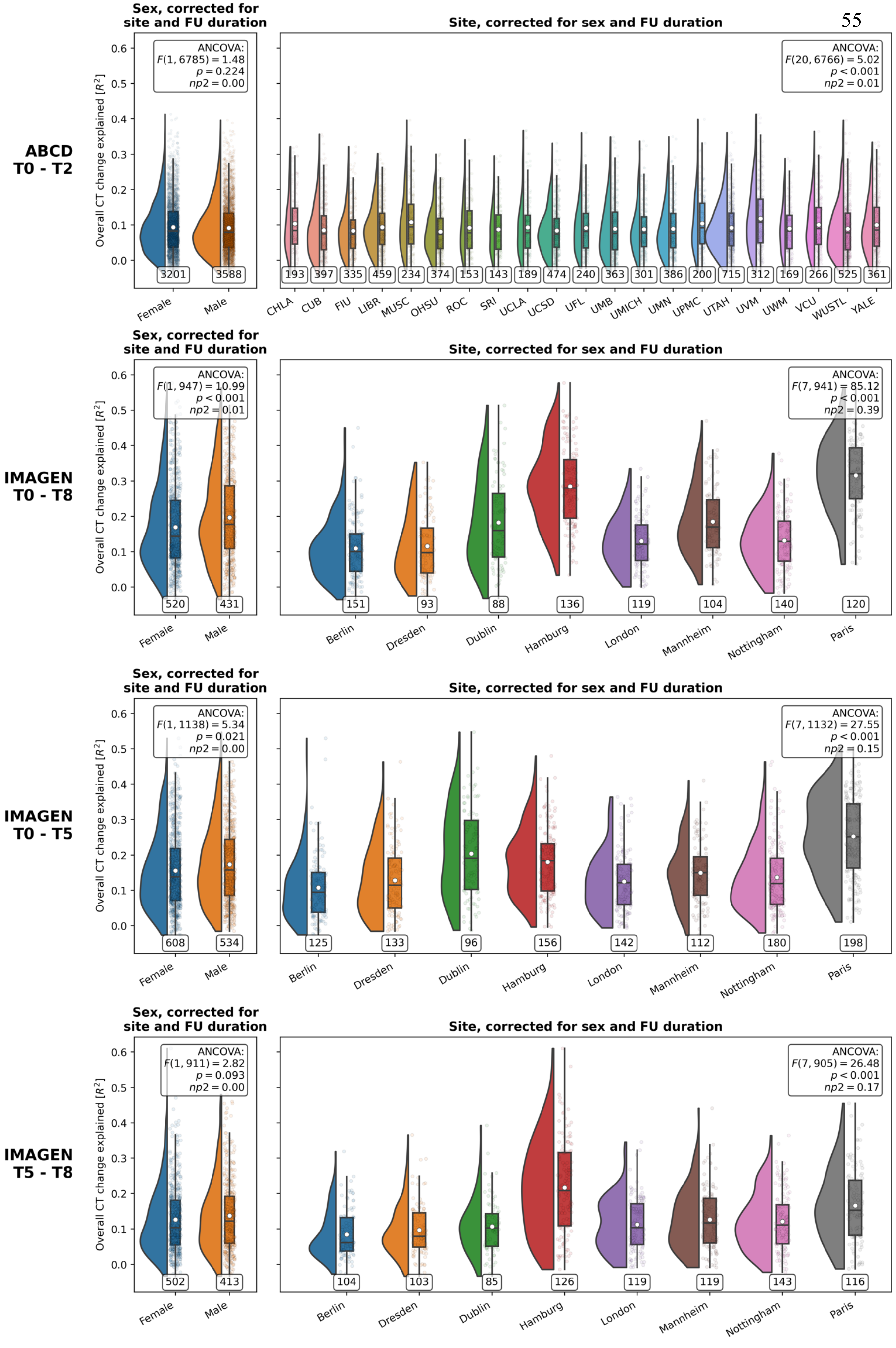
ABCD
T0 - T2
IMAGEN
T0 - T8
IMAGEN
T0 - T5
IMAGEN
T5 - T8
Sex, corrected for site and FU duration
Site, corrected for sex and FU duration
Overall CT change explained [$R^2$]
ANCOVA: $F(1, 6785) = 1.48$ $p = 0.224$ $np2 = 0.00$
ANCOVA: $F(20, 6766) = 5.02$ $p < 0.001$ $np2 = 0.01$
ANCOVA: $F(1, 947) = 10.99$ $p < 0.001$ $np2 = 0.01$
ANCOVA: $F(7, 941) = 85.12$ $p < 0.001$ $np2 = 0.39$
ANCOVA: $F(1, 1138) = 5.34$ $p = 0.021$ $np2 = 0.00$
ANCOVA: $F(7, 1132) = 27.55$ $p < 0.001$ $np2 = 0.15$
ANCOVA: $F(1, 911) = 2.82$ $p = 0.093$ $np2 = 0.00$
ANCOVA: $F(7, 905) = 26.48$ $p < 0.001$ $np2 = 0.17$
Female 3201, Male 3588
CHLA 193, CUB 397, FIU 335, LIBR 459, MUSC 234, OHSU 374, ROC 153, SRI 143, UCLA 189, UCSD 474, UFL 240, UMB 363, UMICH 301, UMN 386, UPMC 200, UTAH 715, UVM 312, UWM 169, VCU 266, WUSTL 525, YALE 361
Female 520, Male 431
Berlin 151, Dresden 93, Dublin 88, Hamburg 136, London 119, Mannheim 104, Nottingham 140, Paris 120
Female 608, Male 534
Berlin 125, Dresden 133, Dublin 96, Hamburg 156, London 142, Mannheim 112, Nottingham 180, Paris 198
Female 502, Male 413
Berlin 104, Dresden 103, Dublin 85, Hamburg 126, London 119, Mannheim 119, Nottingham 143, Paris 116

**Fig. S26: Effects of sex and site on total explained cortical thickness change variance**

Comparisons testing if the overall extent to which ABCD and IMAGEN CT change was explained (**y axes**) across each tested time period (**rows**) varied by binary sex (**column 1**) and site (**column 2**). Note that these analyses are done after site harmonization of the "cross-sectional" timepoint-wise CT data using ComBat. **Raincloud plot** elements as described above. Sex and site were compared using analyses of covariance (ANCOVAs) assessing the effect of (**column 1**) sex on explained CT change variance, including site and follow-up duration as covariates, or (**column 2**) site on explained CT change variance, including binary sex and follow-up duration as covariates. Effect sizes are expressed as eta-squared (np2). Group-level source data are provided as a Source Data file.

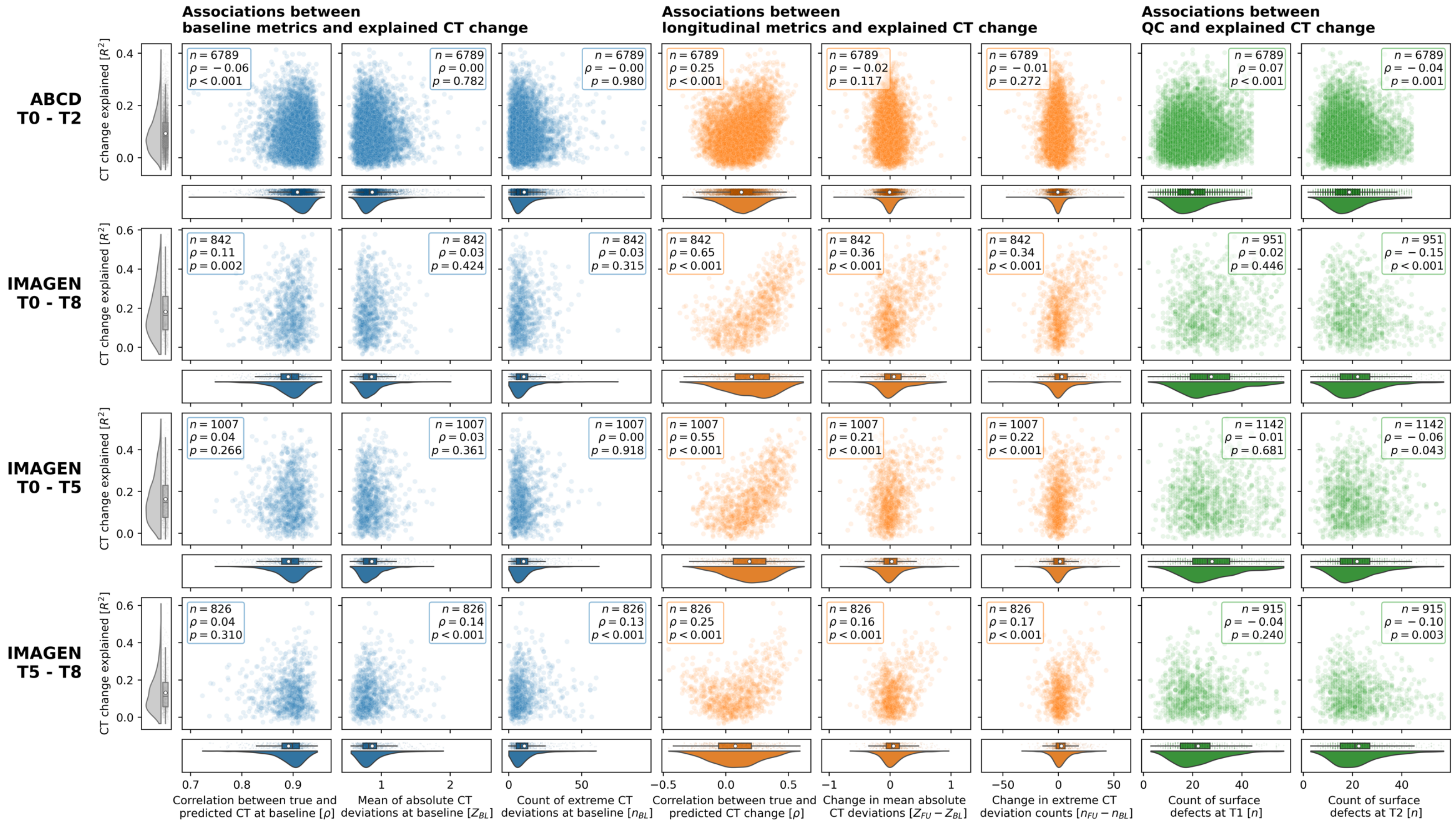

Associations between baseline metrics and explained CT change
Associations between longitudinal metrics and explained CT change
Associations between QC and explained CT change
ABCD T0 - T2
IMAGEN T0 - T8
IMAGEN T0 - T5
IMAGEN T5 - T8
CT change explained [$R^2$]
Correlation between true and predicted CT at baseline [$\rho$]
Mean of absolute CT deviations at baseline [$Z_{BL}$]
Count of extreme CT deviations at baseline [$n_{BL}$]
Correlation between true and predicted CT change [$\rho$]
Change in mean absolute CT deviations [$Z_{FU} - Z_{BL}$]
Change in extreme CT deviation counts [$n_{FU} - n_{BL}$]
Count of surface defects at T1 [$n$]
Count of surface defects at T2 [$n$]

**Fig. S27: Effects of predictive performance, subject-level atypical development, and surface reconstruction quality on explained cortical thickness**

**Scatterplots** show relationships between cohort- and time period-wise (**rows**) total explained CT change (**y axes**) as estimated in subject-wise dominance analyses. **X axes**: **Columns 1 and 4**: Model fit; correlation between observed and predicted baseline CT or CT change. **Columns 2 and 5**: CT deviations; absolute CT deviation Z scores or their longitudinal change. **Columns 3 and 6**: CT deviations; counts of extreme deviations per subject or their longitudinal change (defined as deviation Z > 2). **Columns 7 and 8**: Data quality: Total Euler number, FreeSurfer's quality control metric for surface reconstruction. **Blue plots** indicate baseline metrics, **orange plots** indicate longitudinal metrics, **green plots** indicate associations with the Euler number at the first and second timepoint of each studied time period. **Boxes** in the upper corners of each panel contain Spearman's rho and the associated parametric *p* value for each bivariate association. Group-level source data are provided as a Source Data file.

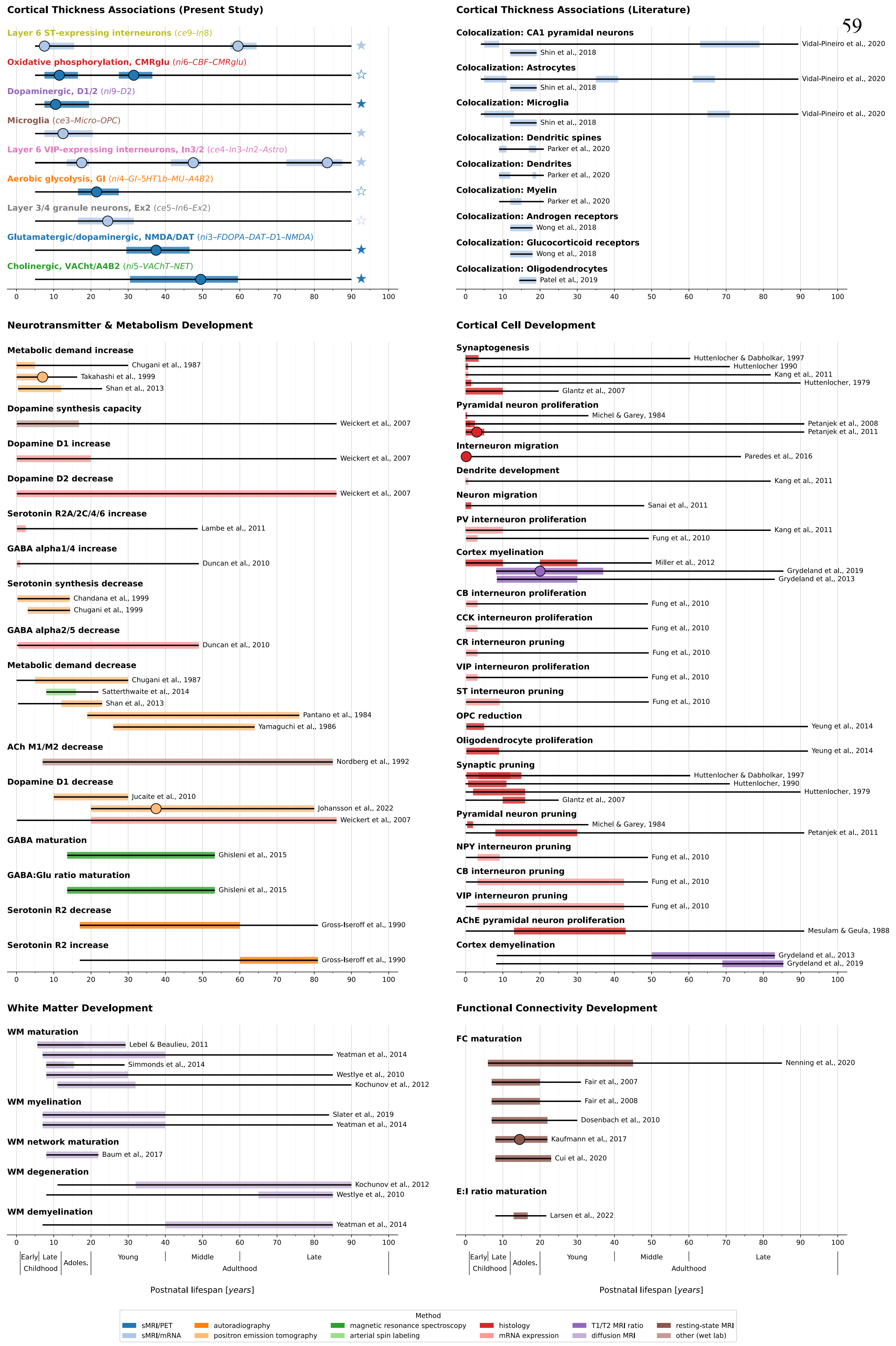
Cortical Thickness Associations (Present Study)
Layer 6 ST-expressing interneurons (ce9–In8)
Oxidative phosphorylation, CMRglu (ni6–CBF–CMRglu)
Dopaminergic, D1/2 (ni9–D2)
Microglia (ce3–Micro–OPC)
Layer 6 VIP-expressing interneurons, In3/2 (ce4–In3–In2–Astro)
Aerobic glycolysis, GI (ni4–GI–5HT1b–MU–A4B2)
Layer 3/4 granule neurons, Ex2 (ce5–In6–Ex2)
Glutamatergic/dopaminergic, NMDA/DAT (ni3–FDOPA–DAT–D1–NMDA)
Cholinergic, VAChT/A4B2 (ni5–VAChT–NET)
Cortical Thickness Associations (Literature)
Colocalization: CA1 pyramidal neurons
Vidal-Pineiro et al., 2020
Shin et al., 2018
Colocalization: Astrocytes
Vidal-Pineiro et al., 2020
Shin et al., 2018
Colocalization: Microglia
Vidal-Pineiro et al., 2020
Shin et al., 2018
Colocalization: Dendritic spines
Parker et al., 2020
Colocalization: Dendrites
Parker et al., 2020
Colocalization: Myelin
Parker et al., 2020
Colocalization: Androgen receptors
Wong et al., 2018
Colocalization: Glucocorticoid receptors
Wong et al., 2018
Colocalization: Oligodendrocytes
Patel et al., 2019
Neurotransmitter & Metabolism Development
Metabolic demand increase
Chugani et al., 1987
Takahashi et al., 1999
Shan et al., 2013
Dopamine synthesis capacity
Weickert et al., 2007
Dopamine D1 increase
Weickert et al., 2007
Dopamine D2 decrease
Weickert et al., 2007
Serotonin R2A/2C/4/6 increase
Lambe et al., 2011
GABA alpha1/4 increase
Duncan et al., 2010
Serotonin synthesis decrease
Chandana et al., 1999
Chugani et al., 1999
GABA alpha2/5 decrease
Duncan et al., 2010
Metabolic demand decrease
Chugani et al., 1987
Satterthwaite et al., 2014
Shan et al., 2013
Pantano et al., 1984
Yamaguchi et al., 1986
ACh M1/M2 decrease
Nordberg et al., 1992
Dopamine D1 decrease
Jucaite et al., 2010
Johansson et al., 2022
Weickert et al., 2007
GABA maturation
Ghisleni et al., 2015
GABA:Glu ratio maturation
Ghisleni et al., 2015
Serotonin R2 decrease
Gross-Iseroff et al., 1990
Serotonin R2 increase
Gross-Iseroff et al., 1990
Cortical Cell Development
Synaptogenesis
Huttenlocher & Dabholkar, 1997
Huttenlocher 1990
Kang et al., 2011
Huttenlocher, 1979
Glantz et al., 2007
Pyramidal neuron proliferation
Michel & Garey, 1984
Petanjek et al., 2008
Petanjek et al., 2011
Interneuron migration
Paredes et al., 2016
Dendrite development
Kang et al., 2011
Neuron migration
Sanai et al., 2011
PV interneuron proliferation
Kang et al., 2011
Fung et al., 2010
Cortex myelination
Miller et al., 2012
Grydeland et al., 2019
Grydeland et al., 2013
CB interneuron proliferation
Fung et al., 2010
CCK interneuron proliferation
Fung et al., 2010
CR interneuron pruning
Fung et al., 2010
VIP interneuron proliferation
Fung et al., 2010
ST interneuron pruning
Fung et al., 2010
OPC reduction
Yeung et al., 2014
Oligodendrocyte proliferation
Yeung et al., 2014
Synaptic pruning
Huttenlocher & Dabholkar, 1997
Huttenlocher, 1990
Huttenlocher, 1979
Glantz et al., 2007
Pyramidal neuron pruning
Michel & Garey, 1984
Petanjek et al., 2011
NPY interneuron pruning
Fung et al., 2010
CB interneuron pruning
Fung et al., 2010
VIP interneuron pruning
Fung et al., 2010
AChE pyramidal neuron proliferation
Mesulam & Geula, 1988
Cortex demyelination
Grydeland et al., 2013
Grydeland et al., 2019
White Matter Development
WM maturation
Lebel & Beaulieu, 2011
Yeatman et al., 2014
Simmonds et al., 2014
Westlye et al., 2010
Kochunov et al., 2012
WM myelination
Slater et al., 2019
Yeatman et al., 2014
WM network maturation
Baum et al., 2017
WM degeneration
Kochunov et al., 2012
Westlye et al., 2010
WM demyelination
Yeatman et al., 2014
Functional Connectivity Development
FC maturation
Nenning et al., 2020
Fair et al., 2007
Fair et al., 2008
Dosenbach et al., 2010
Kaufmann et al., 2017
Cui et al., 2020
E:I ratio maturation
Larsen et al., 2022
0 10 20 30 40 50 60 70 80 90 100
Early Childhood
Late Childhood
Adoles.
Young
Middle
Late
Adulthood
Postnatal lifespan [years]
Method
sMRI/PET
sMRI/mRNA
autoradiography
positron emission tomography
magnetic resonance spectroscopy
arterial spin labeling
histology
mRNA expression
T1/T2 MRI ratio
diffusion MRI
resting-state MRI
other (wet lab)

**Fig. S28: Summary of previous literature on processes involved in human brain development and maturation**

Visualization of published results on correlates of human CT development and maturation, extention to Fig. 10. This collection is not exhaustive and is mostly limited to *in vivo* and *postmortem* studies in humans. The **upper left** panel shows results drom the current study. **Thin black lines** depict the age range covered by each study. **Thick colored bars** show the time period in which the respective study target was reported to show developmental changes. If available, peaks of these assocations are marked by **circles**. Results are **grouped** by the broad area of research, **colors** code the applied methodology. All references can be found in the supplementary reference list[55–75,75–82,82–105]. Source data are provided as a Source Data file.
