## Supplementary figures and images for "Regional patterns of human cortex development correlate with underlying neurobiology"

### Supplementary Animation S1

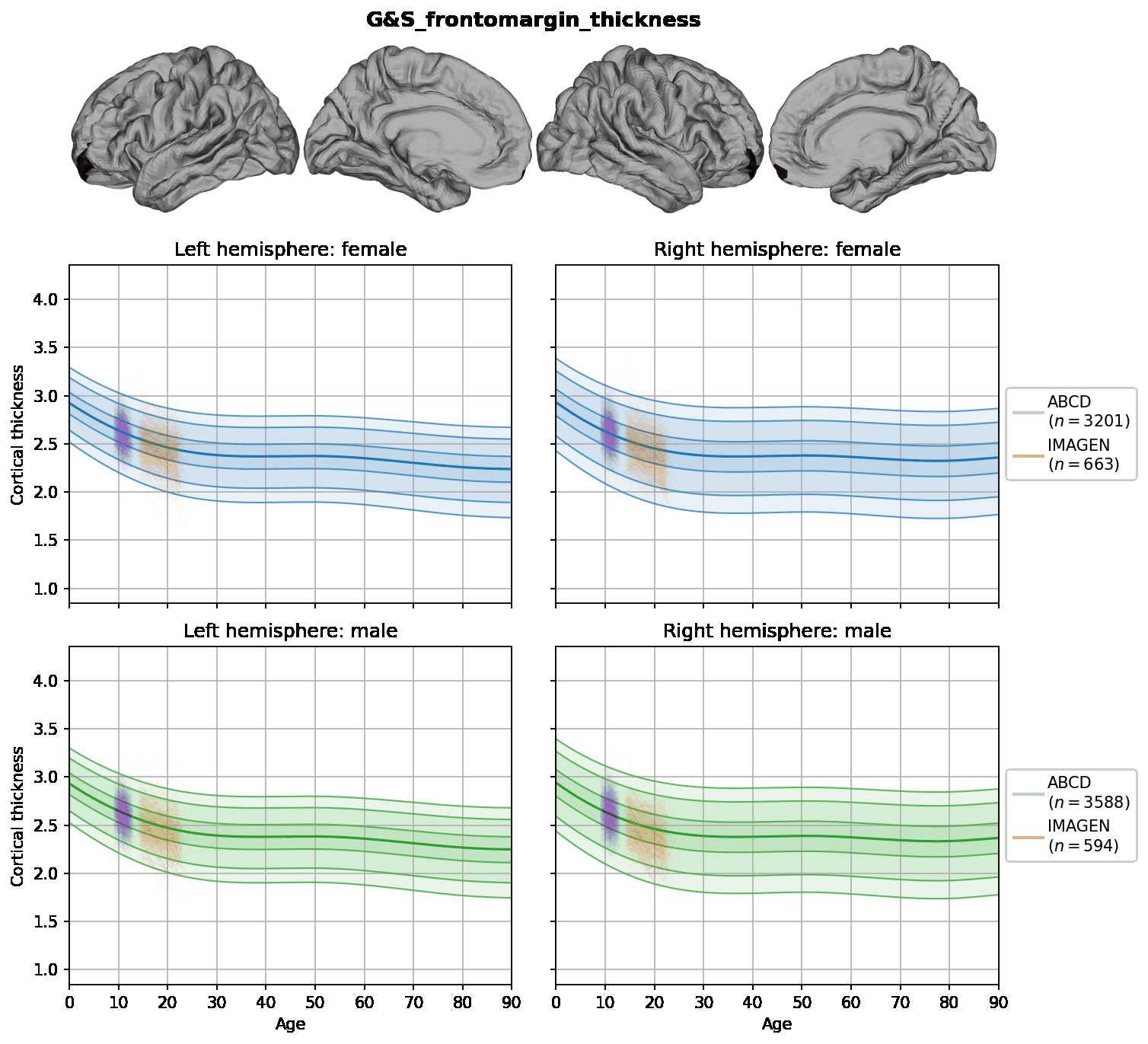

### Supplementary Animation S2

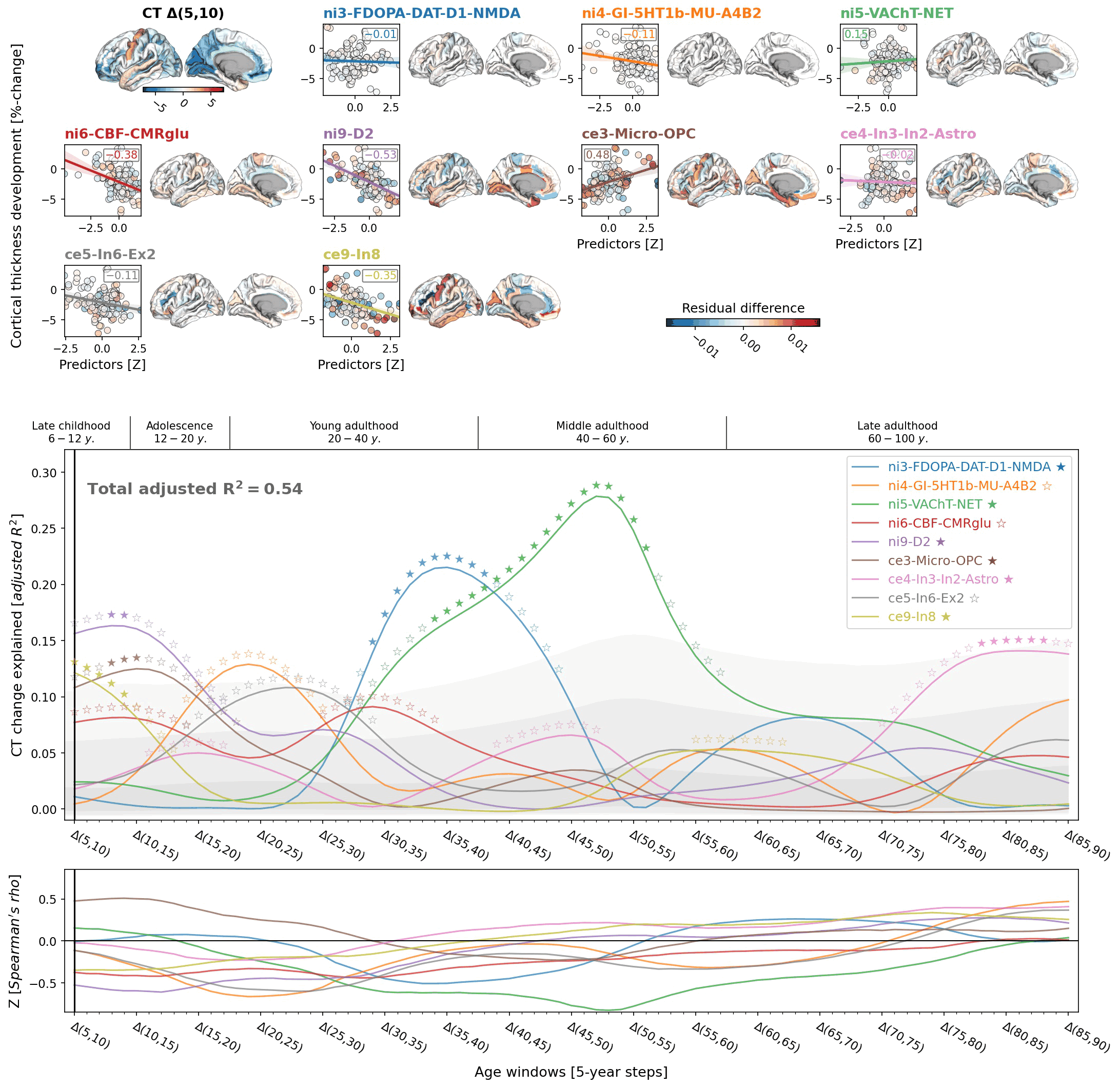
